# Glial place cells: complementary encoding of spatial information in hippocampal astrocytes

**DOI:** 10.1101/2021.07.06.451296

**Authors:** Sebastiano Curreli, Jacopo Bonato, Sara Romanzi, Stefano Panzeri, Tommaso Fellin

## Abstract

Calcium dynamics into astrocytes influence the activity of nearby neuronal structures. However, because previous reports show that astrocytic calcium signals largely mirror neighboring neuronal activity, current information coding models neglect astrocytes. Using simultaneous two-photon calcium imaging of astrocytes and neurons in the hippocampus of mice navigating a virtual environment, we demonstrate that astrocytic calcium signals actively encode spatial information. Calcium events carrying spatial information occurred in topographically organized astrocytic subregions. Importantly, astrocytes encoded spatial information that was complementary and synergistic to that carried by neurons, improving spatial position decoding when astrocytic signals were considered alongside neuronal ones. These results suggest that the complementary place-dependence of localized astrocytic calcium signals regulates clusters of nearby synapses, enabling dynamic, context-dependent, variations in population coding within brain circuits.

## Main Text

Astrocytes, the most abundant class of glial cells in the brain, exhibit complex dynamics in intracellular calcium concentration ^1^. Intracellular calcium signals can be spatially restricted to individual subcellular domains (e.g., cellular processes *vs* somata) and be coordinated across astrocytic cells ^2–8^. In the intact brain, astrocytic calcium dynamics can be spontaneous ^9^ or triggered by the presentation of external physical stimuli ^4, 7, 10–12^. Interestingly, previous reports suggest that astrocytic calcium signals triggered by external sensory stimuli largely mirror the activity of local neuronal cells ^10, 11^. Such findings have led current models of sensory information coding in the brain to overlook the contribution of astrocytes, under the implicit or explicit assumption that astrocytic cells only provide information already encoded in neurons ^13, 14^. Here, we challenged this assumption and tested the hypothesis that astrocytes encode information in their intracellular calcium dynamics that is not present in the activity of nearby neurons. As a model, we used spatial information encoding in the hippocampus, where neural place cells encode navigational information by modulating their firing rate as a function of the animal’s spatial location ^15–17^. We demonstrate that astrocytic calcium signals actively encode information about the animal’s position in virtual space, and that this information is complementary to that carried by hippocampal neurons.

We combined two-photon functional imaging in head-fixed mice navigating in virtual reality ^16, 17^ (Fig. 1A) with astrocyte-specific expression of the genetically encoded calcium indicator GCaMP6f (Fig. 1B, D) ^18–20^. We measured subcellular calcium dynamics of hippocampal CA1 astrocytes during spatial navigation in a virtual monodirectional corridor (Fig. 1C) ^21^. Using the intersection of two stringent criteria (significance of mutual information about spatial location carried by the cell’s activity, and reliability of calcium activity across running trials; Methods, Extended Data Fig 1), we found that a large fraction of astrocytic regions of interest (ROIs) had calcium signals that were reliably modulated by the spatial position of the animal in the virtual track (44 ± 21 %, 155 out of 356 ROIs, from 7 imaging sessions on 3 animals, Fig. 1E, Extended Data Table 1). We defined the spatial response field of an astrocytic ROI as the portion of virtual corridor at which that ROI showed, on average across trials, increased GCaMP6f fluorescence (Methods). The distribution of astrocytic spatial response field positions covered the entire length of the virtual corridor (Fig. 1F, G, N = 155 ROIs from 7 imaging sessions on 3 animals). The median width of the astrocytic spatial field was 56 ± 22 cm (N = 155 ROIs from 7 imaging sessions in 3 animals, Fig. 1H). ROIs with reliable spatial information had reproducible estimates of spatial response profiles (Extended Data Fig. 2). Splitting the dataset in odd and even trials resulted in a similar distribution of astrocytic field position compared to the entire dataset (Fig. 1F center and rightmost panels, Fig. 1I). Experiments performed with mice trained in a bidirectional virtual environment (Extended Data Fig. 3) ^16, 17^ confirmed the results obtained in the monodirectional virtual environment: a significant fraction of astrocytic ROIs carried significant information about the spatial position of the animal in the virtual corridor and the distribution of position of the astrocytic spatial field covered the whole virtual corridor (29 ± 13 %, N = 192 out of 648 ROIs in the forward direction; 20 ± 13 %, N = 133 out of 648 ROIs in the backward direction, p = 0.09 Wilcoxon signed rank test for comparison between forward and backward directions, from 18 imaging sessions in 4 animals; Extended Data Fig. 3E, F). The median width of the spatial response field was 44 ± 20 cm, N = 192 out of 648 ROIs in the forward direction and 44 ± 29 cm, N = 133 out of 648 ROIs in the backward direction (p = 0.34 Wilcoxon rank-sums test for comparison between forward and backward directions, Extended Data Fig. 3G). In the bidirectional environment, astrocytic ROIs showed significant direction-selective spatial modulation in their response field (Extended Data Fig. 3H). Thus, astrocytic calcium signaling participates in encoding spatial information in the hippocampus.

**Figure 1.**
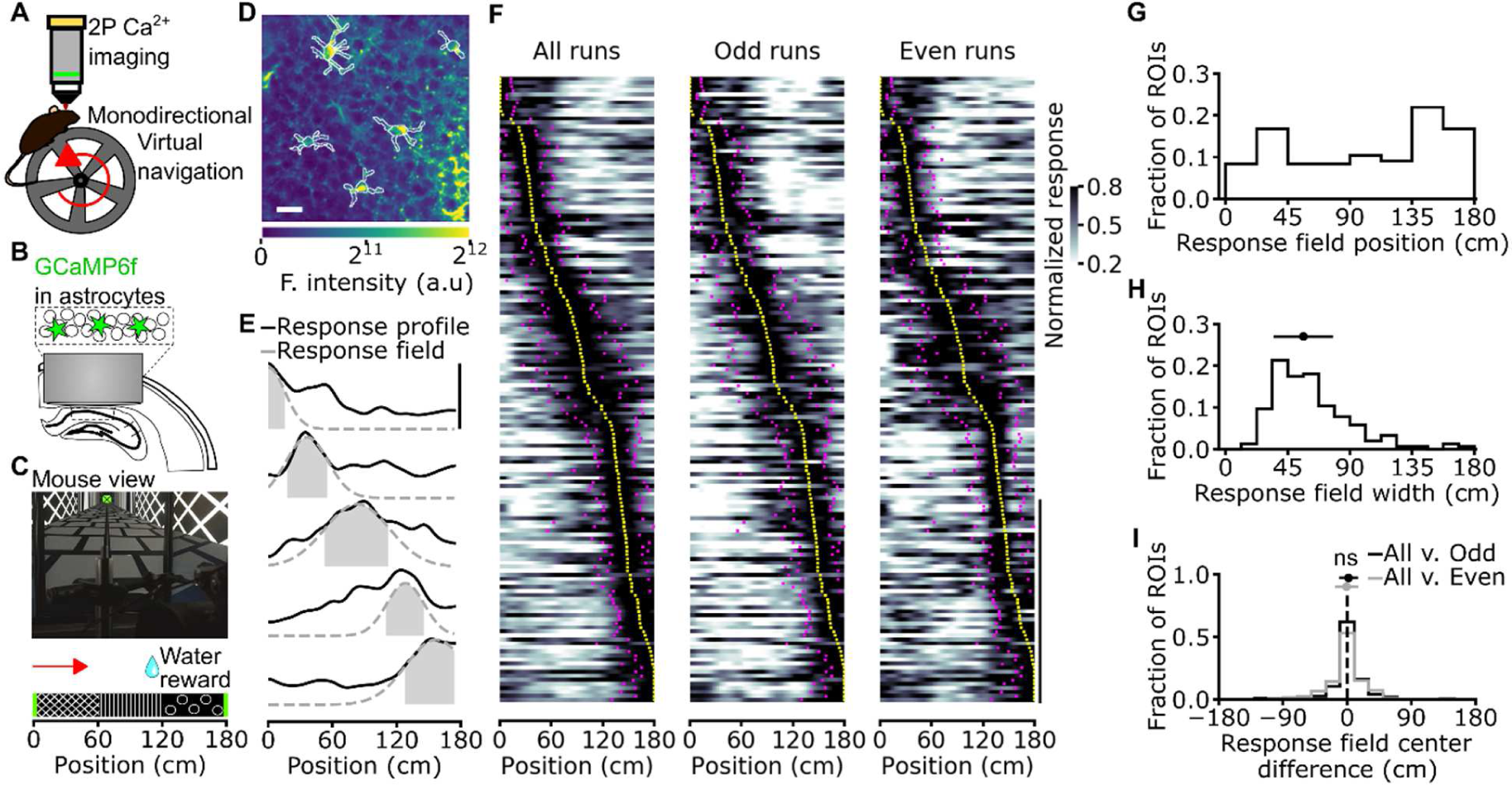
Astrocytic calcium signals in the CA1 hippocampal area encode spatial information during virtual navigation. (A) Two-photon fluorescence imaging was performed in head-fixed mice running along a monodirectional virtual track. (B) GCaMP6f was expressed in CA1 astrocytes and imaging was performed through a chronic optical window. (C) Mice navigated in a virtual linear corridor in one direction, receiving a water reward in the second half of the virtual corridor. (D) Median projection of GCaMP6f-labeled astrocytes in the CA1 pyramidal layer. Scale bar: 20 µm. (E) Calcium signals for five representative astrocytic ROIs encoding spatial information across the corridor length. Solid black lines indicate the average astrocytic calcium response across trials as a function of spatial position. Dashed grey lines and filled grey areas indicate Gaussian fitting function and response field width (see Methods), respectively. (F) Normalized astrocytic calcium responses as a function of position for astrocytic ROIs that contain significant spatial information (n = 155 ROIs with reliable spatial information out of 356 total ROIs, 7 imaging sessions from 3 animals). Responses are ordered according to the position of the center of the response field (from minimum to maximum). Left panel, astrocytic calcium responses from all trials. Center and right panels, astrocytic calcium responses from odd (center) or even (right) trials. Yellow dots indicate the center position of the response field, magenta dots indicate the extension of the field response (see Methods, vertical scale: 50 ROIs). (G) Distribution of response field position. (H) Distribution of field width. (I) Distribution of the differences between the center position of the response fields in cross-validated trials and odd trials (black) or cross-validated and even trails (grey). Deviations for odd and even trials are centered at 0 cm: median deviation for odd trials 2 ± 13 cm; median deviation for even trials -1 ± 17 cm, neither is significantly different from zero (p = 0.07 and p = 0.69, respectively, Wilcoxon signed-rank test with Bonferroni correction. N = 155 ROIs from 7 imaging sessions on 3 animals).

**Figure 2.**
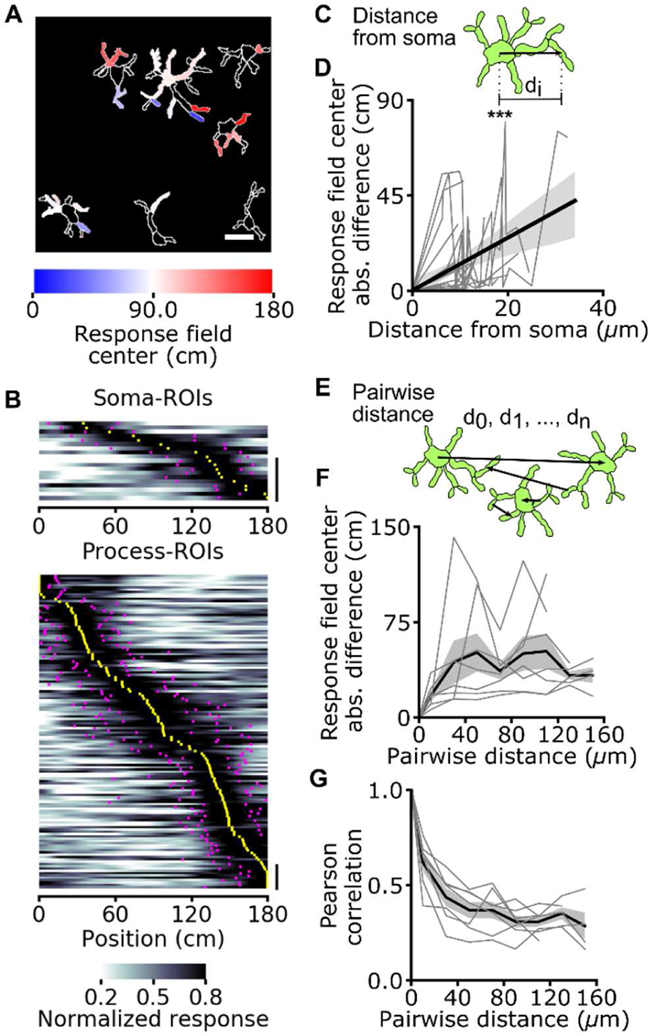
Topographic organization of spatial information encoding in astrocytes: somas *vs* processes. (A) Astrocytic ROIs in a representative FOV are color-coded according to response field position along the virtual corridor. Scale 20 µ m. (B) Normalized astrocytic calcium responses as a function of position for astrocytic ROIs with reliable spatial information corresponding to somas (top) and processes (bottom) (somas: 19 ROIs with reliable spatial information out of 46 total ROIs; processes: 136 ROIs with reliable spatial information out of 310 total ROIs; data from 7 imaging sessions in 3 animals). Vertical scale: 10 ROIs. (C) Distance between the center of a process-ROI and corresponding soma-ROI computed for each astrocyte. (D) Absolute difference in response field position of a process-ROI with respect to the field position of the corresponding soma-ROI as a function of the distance between the two (R^2^ = 0.21, p = 3.2E-6, Wald test, data from 19 cells in which there was significant spatial modulation in the soma and at least one process; 7 imaging sessions on 3 animals). (E) The distance between the centers of pairs of ROIs (d_0_, d_1_, d_n_) is computed across recorded astrocytic ROIs. (F, G) Pearson’s correlation (F) and difference between response field position (G) for pairs of astrocytic ROIs containing reliable spatial information across the whole FOV as a function of pairwise ROI distance. Grey lines indicate single experiments, black line and the grey shade indicate mean ± s.e.m, respectively. Data from 7 imaging sessions in 3 animals. *, p < 0.05; **, p ≤ 0.01; ***, p ≤ 0.001.

**Figure 3.**
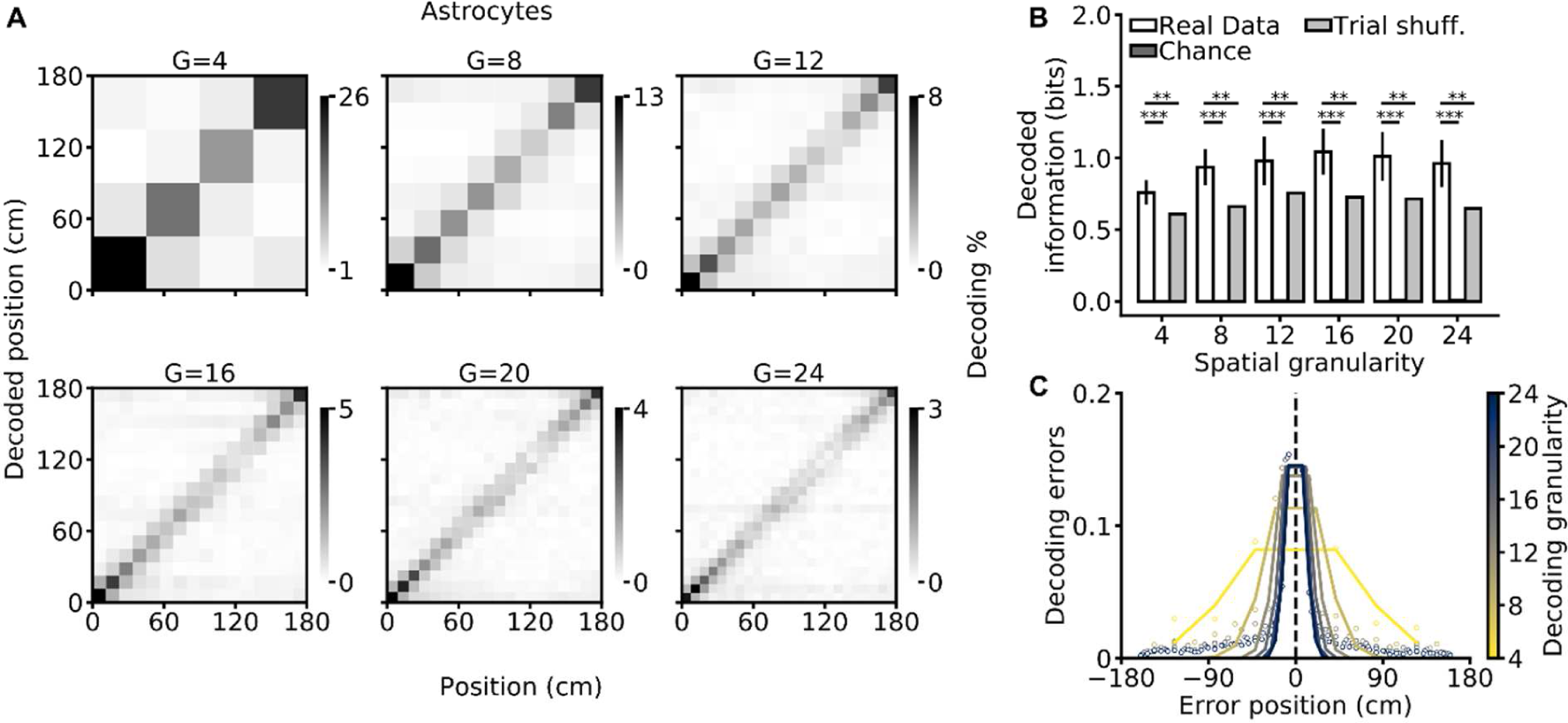
Efficient decoding of the animal’s spatial location from astrocytic calcium signals. (A) Confusion matrices of a SVM classifier for different decoding granularities (G = 4, 8, 12, 16, 20, 24). The actual position of the animal is shown on the x-axis, decoded position is on the y-axis. The grey scale indicates the number of events in each matrix element. (B) Decoded information as a function of decoding granularity on real (white), chance (dark gray), and trial-shuffled (grey) data (see methods). Trial-shuffling disrupts temporal coupling within astrocytic population vectors while preserving single-ROI activity patterns. Data are shown as mean ± s.e.m. See also Extended Data Table 2. (C) Decoding error as a function of the error position within the confusion matrix. The color code indicates decoding granularity. Data in all panels were obtained from 7 imaging sessions in 3 animals.

Astrocytic calcium signaling has been shown to be organized at the subcellular level; the calcium dynamics of astrocytic cellular processes can be distinct from those occurring in the astrocytic cell body ^2, 3, 5, 7, 8^. We thus categorized astrocytic ROIs (among the set of 356 described above) according to whether they were located within main processes (process-ROIs) or cell bodies (soma-ROIs, Fig. 2). Signals from both soma-ROIs and process-ROIs encoded spatial information (Fig. 2A). Moreover, a similar fraction of soma-ROIs and process-ROIs were modulated by the spatial position of the animal (42 ± 34 %, 19 out of 46 soma-ROIs *vs* 44 ± 21 %, 136 out of 310 process-ROIs, p = 0.61 Wilcoxon signed-rank test, from 7 imaging sessions on 3 animals). The distribution of field position of soma-ROIs and process-ROIs similarly covered the entire length of the virtual corridor (Fig. 2B, Extended Data Fig. 4A, Table 1). The average width of the astrocytic spatial field was not statistically different between process-ROIs and soma-ROIs (Extended Data Fig. 4B). Within individual astrocytes, the difference between the field position of a process-ROI and the corresponding soma-ROI (both containing reliable spatial information) increased as a function of the distance between the two ROIs (Fig. 2C, Extended Data Fig. 4). Thus, spatial information was differentially encoded in topographically distinct locations of the same astrocyte. The difference between the field position of a process-ROI and the corresponding soma-ROI did not depend on the angular position of the process with respect to the soma (Extended Data Fig. 4). When comparing calcium activity across pairs of ROIs with reliable spatial information (belonging to processes or somas across astrocytes), correlation decreased as a function of the pair distance (τ_decay_ = 14 ± 2 µm, R^2^ = 0.98) in the 0-50 µm range, and then substantially plateaued for pair distances between 50 µm and 160 µm (Fig. 2F). This indicates that calcium signals encoding reliable spatial information were coordinated across distant ROIs, even those putatively belonging to different cells. In agreement with this observation, the difference in field position among pairs of ROIs with reliable spatial information increased as a function of pair distance within 0-40 µ m and then plateaued to a constant value (τ_rise_ = 13 ± 7 µm, R^2^ = 0.79) for pair distances between 40-160 µ m (Fig. 2G). Event-triggered averages of astrocytic responses representing temporal relationships between calcium signals at different subcellular regions are shown in Extended Data Fig. 5.

**Figure 4.**
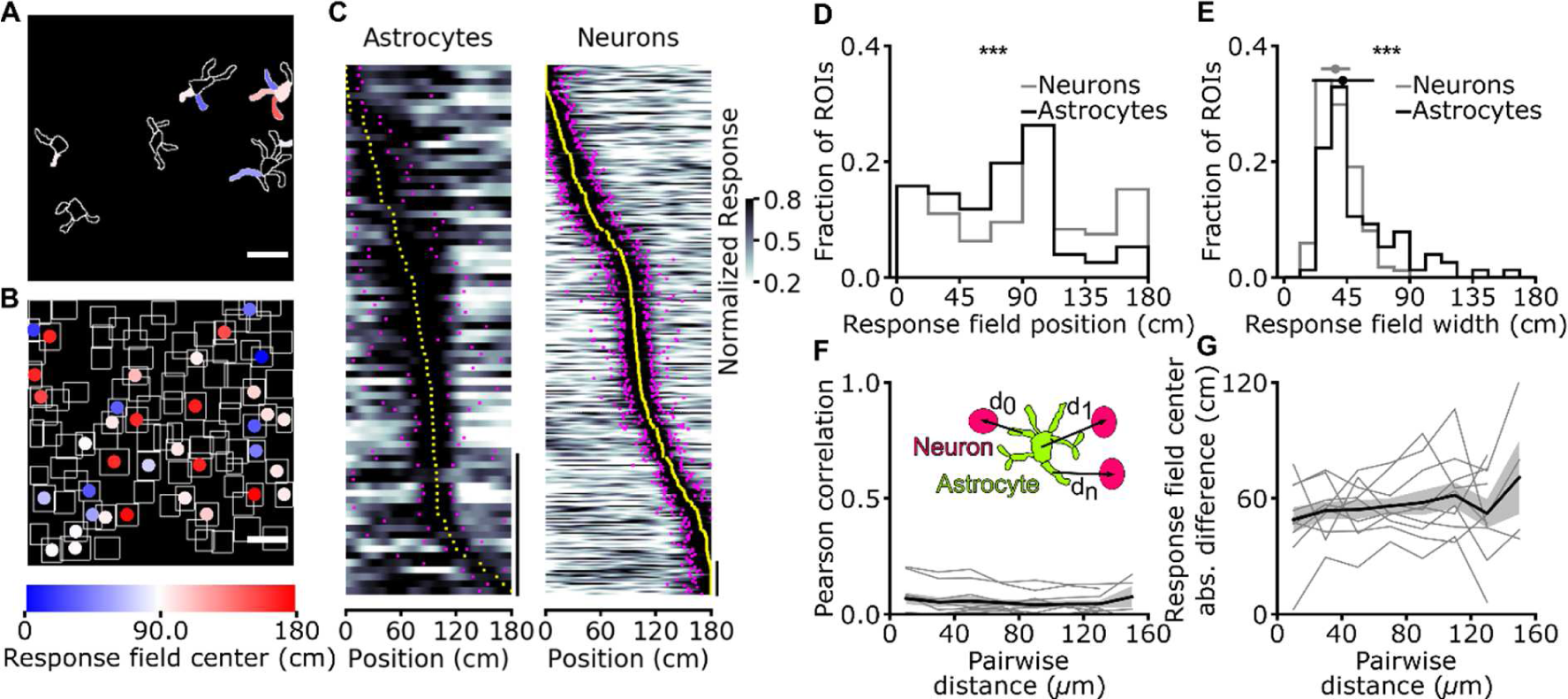
Astrocytes have broader response field width and a different distribution of field position compared to neurons. (A, B) ROIs corresponding to simultaneously recorded GCaMP6f-labeled astrocytes (A) and jRCaMP1a-labeled neurons (B) in the CA1 pyramidal layer. ROIs are color-coded according to response field and place field center along the virtual corridor, respectively. Scale bar, 20 µ m. (C) Normalized calcium responses as a function of position for astrocytic ROIs (left) and neuronal ROIs (right) that contain a significant amount of spatial information (astrocytic ROIs, N = 76 ROIs with reliable spatial information out of 341 total ROIs; neuronal ROIs, N = 335 ROIs with reliable spatial information out of 870 total ROIs, data from 11 imaging sessions in 7 animals). Responses are ordered according to the position of the center of the response field for astrocytes and place field for neurons. Vertical scale bar, 20 ROIs. (D) Distribution of astrocytic response field position (black line) and neuronal place field position (grey line, p = 5E-4, Kolmogorov-Smirnov test for comparison between astrocytic and neuronal distribution). (E) Distribution of astrocytic response field width (black line) and neuronal place field width (grey line, median width of astrocytic response field: 42 ± 22 cm, N = 76; median width of neuronal place field: 37 ± 10 cm, N = 335, p = 2E-5, Wilcoxon Rank-sum test for comparison between astrocytic and neuronal distribution). (F, G) The inset shows astrocytic ROIs (green) and neuronal ROIs (pink). For all pairs, the distance (d_0_, d_1_, d_n_) between the center of an astrocytic ROI and the center of a neuronal ROI, both containing reliable spatial information, is computed. Pairwise Pearson’s correlation (F) and difference between response field position for astrocyte-neuron ROI pairs (G) as a function of pair distance. Data are from 11 imaging sessions in 7 animals (see also Extended Data Fig. 10).

**Figure 5.**
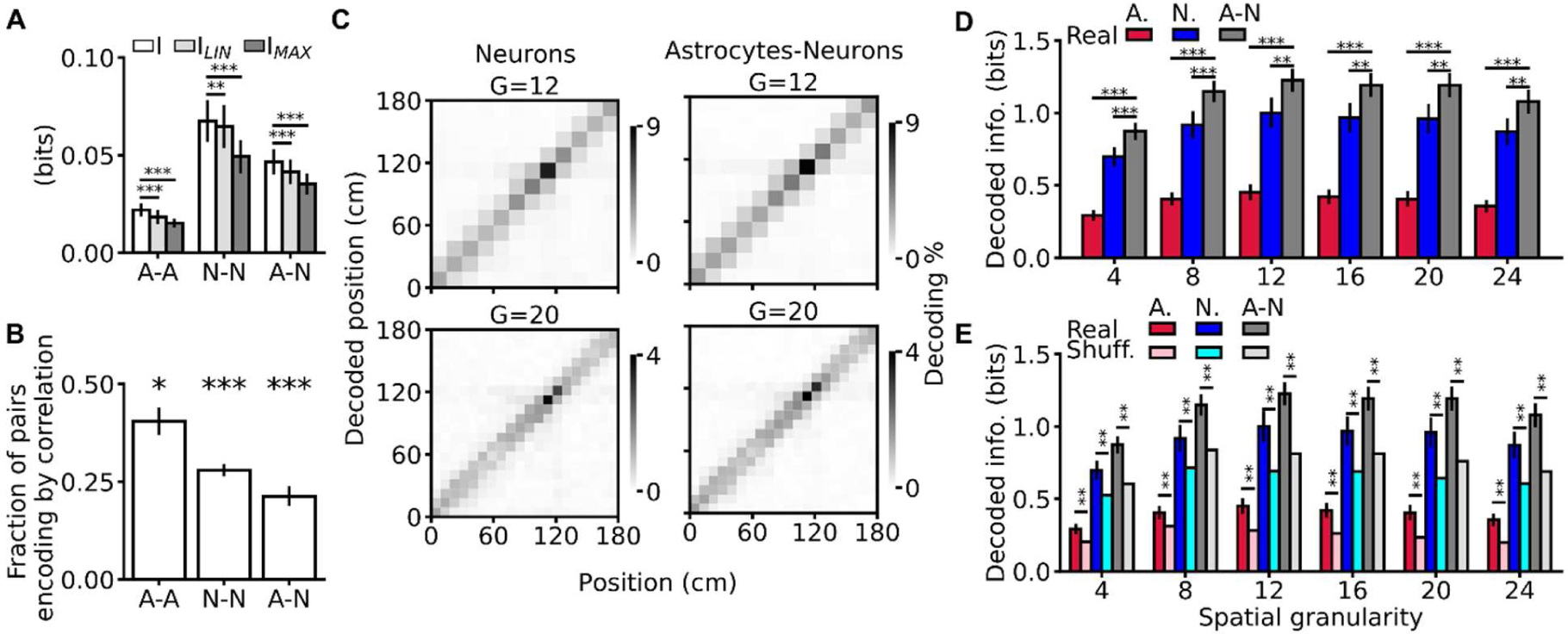
Spatial information encoding in astrocytes is complementary and synergistic to spatial information encoding in neurons. (A) Information about position carried by pairs of ROIs (I) compared the sum (I_LIN_) or the maximum (I_MAX_) of the information separately encoded by each member of the pair. A-A, pair composed of two astrocytic ROIs; N-N, pair composed of two neuronal ROIs; A-N, mixed pair composed of one astrocytic and one neuronal ROI (I vs. I_LIN_: A-A: p = 1E-3, N-N: p = 5E-3, A-N: p = 1E-3; I vs. I_MAX_: A-A: p = 1E-3, N-N: p = 1E-3, A-N: p = 1E-3, Wilcoxon signed-rank test). (B) Fraction of pairs encoding spatial information encoding by correlations (A-A: p = 3E-2, N-N: p = 1E-3, A-N: p = 1E-3, Wilcoxon signed rank-test with respect to the null hypothesis that a pair could be either synergistic or non-synergistic with equal probability set at 0.5). (C) Representative confusion matrices of a SVM classifier decoding mouse position using population vectors comprising neuronal (left) or astrocytic and neuronal ROIs (right), for different decoding granularities (G = 12, 20, see also Extended Data Fig. 13). (D) Decoded information for population vectors of different compositions (A, astrocytic ROIs only; N, neuronal ROIs only; A-N, population vector considering all ROIs) as a function of decoding granularity (see Extended Data Table 3). (E) Same as in (D) but adding comparison with trial-shuffled data (lighter bars) (see Extended Data Table 4). In panels A-B, D-E data are represented as mean ± s.e.m. In all panels, data are obtained from 11 imaging sessions in 7 animals.

Since calcium dynamics of individual astrocytic ROIs encodes significant spatial information, it should be possible to decode the animal’s position in the virtual corridor from single-trial calcium dynamics of populations of astrocytic ROIs. We trained a support vector machine (Methods) to classify the mouse’s position according to a set of discrete spatial locations using a single-trial population vector made combining calcium signals of all individual astrocytic ROIs within the FOV. We computed the population decoding accuracy and the decoded spatial information ^22^ as a function of spatial granularity, i.e., the number of discrete locations available to the SVM decoder (4, 8, 12, 16, 20, or 24 locations). We found that the SVM predicted the animal’s spatial location across granularities (Fig. 3A, Extended Data Table 1). Cross-validated decoding accuracy (Extended Data Fig. 6) and decoded spatial information were significantly above chance (Fig. 3B) across the entire range of spatial granularities (chance level was estimated by decoding position after randomly shuffling spatial locations in the data while preserving the temporal structure of the population calcium signals, see Methods). Disrupting the within-trial temporal coupling within astrocytic population vectors while preserving single-ROI activity patterns ^23, 24^ consistently decreased decoded spatial information (Fig. 3B) and decoding accuracy (Extended Data Fig. 6). This suggests that within-trial interactions among astrocytic ROIs encode spatial information not present in their individual activities. Misclassifications were more likely to happen among nearby locations across all granularity conditions (Fig. 3C), consistent with the idea that astrocytic activity allows localization of the animal’s position. Experiments performed with mice trained in a bidirectional virtual environment (Extended Data Fig. 7) largely confirmed these decoding results.

How does the astrocytic representation of spatial information relate to that of neuronal cells? We combined astrocyte-specific expression of GCaMP6f with neuronal expression of jRCaMP1a ^25^ and performed simultaneous dual color hippocampal imaging with two-photon microscopy (Fig. 4A, B) during virtual navigation. We found that a sizable fraction of astrocytic and neuronal ROIs (astrocytes, 22 ± 19 %, 76 out of 341 ROIs; neurons, 38 ± 13 %, 335 out of 870 ROIs, from 11 imaging sessions on 7 animals) reliably encoded information about the spatial position of the animal in the virtual corridor. For both astrocytes and neurons, the distribution of field position covered the entire length of the virtual corridor (Fig. 4C, D). However, the median width of the astrocytic spatial field was larger than that of neurons (Fig. 4E, Extended Data Table 1). Event triggered averages of astrocytic ROIs signals triggered by neuronal signals are shown in Extended Data Fig. 8. We then investigated the organization of astrocytic and neuronal spatial representations across the FOV. We found that calcium dynamics among mixed pairs of ROIs (one astrocytic ROI with reliable spatial information and one neuronal ROI with reliable spatial information) were significantly correlated (Extended Data Fig. 9), independent of pair distance (0-160 µ m; Fig. 4F). Correlation among pairs of astrocytic ROIs was generally higher than correlation among pairs of neuronal ROIs (Extended Data Fig. 9, 10). The difference in spatial field position of an astrocytic ROI with reliable spatial information and a neuronal ROI with reliable spatial information was also largely independent of pair distance (Fig. 4G). The distinct features of spatial representations in neuronal and astrocytic networks and the evidence of interactions among different cells and cell types described above suggest that calcium dynamics in astrocytes and neurons might carry complementary information about space, i.e. that the information carried by the combined astrocytic and neuronal signals may be greater than information carried by either signal alone.

We quantitatively tested this hypothesis at the pairwise level using mutual information analysis ^22^ on all pairs of ROIs (either astrocytic, neuronal, or mixed pairs). Regardless of pair identity, we found that information carried by pairs of ROIs was greater than information carried by either ROI individually (Fig. 5A). Moreover, information carried by pairs of ROIs was higher than the sum of the information carried by each of two ROIs, regardless of pair identity (Fig. 5A, Extended Data Table 1). Thus, information carried by the pairs was also synergistic. To understand how correlations between ROIs leads to synergistic coding, we used mutual information breakdown analysis of ROI pairs ^26, 27^. This revealed two notable results. First, the “signal-similarity” component of information (I_SS_), which quantifies the reduction of ROI pair information, or redundancy, due to the similarity of the trial-averaged response profiles of the individual ROIs (see Methods and Extended Data Fig. 11), was close to zero. Thus, the diversity of spatial profiles allowed ROIs to sum up their information with essentially no redundancy. Second, synergy between elements of pairs was based on a positive stimulus-dependent correlation component (I_CD_, see Methods and Extended Data Fig. 11), which contributed to increase the joint information. Mathematically, I_CD_ can be non-zero if and only if within-trial correlations between ROIs are modulated by the animal’s position and they carry information complementary to that given by position modulation of each individual ROI ^27^. Correlation enhancement of spatial information was found in a sizeable fraction of pairs across all pair identities, including mixed pairs (Fig. 5B). This was because the strength of correlations between neurons and astrocytes marked the position in virtual corridor: for pairs of one neuronal ROI and one astrocytic ROI, the absolute magnitude of correlations showed a position-dependent modulation (Extended Data Fig. 12), with stronger correlations inside the spatial fields.

Complementary and synergistic spatial information encoding in mixed pairs suggests that the astrocytic network carries additional information unique from that encoded in neuronal circuits also at the whole population level. To directly address this hypothesis, we computed the spatial information gained by decoding the animals’ position from an SVM operating on population vectors comprising either all neuronal, all astrocytic, or all ROIs of both types. We found that neuronal, astrocytic, and mixed population vectors allowed to classify the animal’s position across granularity conditions (Fig 5C-E and Extended Data Fig. 13). However, decoding population vectors comprising both astrocytic and neuronal ROIs led to a greater amount of spatial information than decoding either neuronal or astrocytic population vectors separately (Fig 5D). This result proved that the population of astrocytic ROIs carries information not found in neurons or their interactions. In agreement with what we found in the pair analysis, information decoded from all types of population vectors decreased when within-trial temporal correlations between cells were disrupted by trial shuffling (Fig. 5E, Extended Data Fig. 13) ^23, 27^. Within-trial correlations were thus an important factor for the complementary and synergistic contribution of astrocytes to spatial information encoding at the population level.

## Discussion

Our findings demonstrate, for the first time, that information-encoding cellular signals during virtual spatial cognition extend beyond neuronal circuits to include the nearby astrocytic network. This information was expressed in spatially-restricted subcellular regions, including cellular processes and somas, in agreement with previous work describing the complexity and compartmentalization of calcium signals in these glial cells ^2–5, 8, 28^. Importantly, individual astrocytes could encode multiple spatial fields across different subcellular compartments, suggesting that a single astrocyte may integrate multiple neuronal spatial representations. Interestingly, the spatial representations in individual astrocytes displayed a concentric organization: the difference between the place field position of a subcellular process and the place field position of the corresponding soma increases as a function of distance.

Most importantly, the spatial information encoded in the astrocytic and neuronal networks is distinct. First, the width of the astrocytic spatial field was larger than that of neurons, which may be due to astrocytes integrating multiple neuronal spatial fields or to a prevalence of slower calcium dynamics in astrocytes compared to neurons (^29, 30^, but see ^2, 3, 5, 7^). Second, response field position was differentially distributed in astrocytes compared to neurons, suggesting that CA1 astrocytes do not merely mirror position information encoded in CA1 pyramidal neurons. Third, combining astrocytic and neuronal signals generated significantly greater information about the animal’s position, indicating the signals are both complementary and synergistic. The complementary and synergistic information of astrocytes relied both on the diversity of position tuning and on position-dependent correlations among astrocytic and neuronal ROIs similarly to what observed on neuronal ROIs by ^24^. It should also be considered that astrocytic signals may convey complementary information by simultaneously integrating the activity of several different neuronal inputs encoding distinct stimulus-related variables ^31–33^.

Models of hippocampal function posit that information about variables of the external environment, which are key to spatial navigation and memory, is exclusively encoded in population of neurons ^34–37^. Our results challenge this established view by revealing a fundamental new level of organization for information encoding in the hippocampus during virtual navigation: spatial information, not available in the activity of CA1 projecting neuron or in their interactions, is encoded in the calcium dynamics of local non-neuronal elements and in their position-dependent interaction with neurons. The presence of this additional non-neural reservoir of information and the dependence of the interaction between neuron and astrocytes on key cognitive variables reveal novel and unexpected cellular mechanisms underlying how brain circuits encode information.

Can complementary spatial information encoded in astrocytic calcium dynamics contribute to neuronal computation? If so, how? Although our data do not directly address these questions, previous work in other brain regions reported that astrocytic calcium dynamics largely mirror the activity of nearby neurons ^7, 10, 11^ and that astrocytic signals translate into changes in neuronal excitability and synaptic transmission through various mechanisms (reviewed in ^1, 38–40^). In this scenario, changes in synaptic transmission and neuronal excitability induced by astrocytic calcium dynamics that simply mirror neuronal information would only modulate neural activity *without providing further information*, as all the activity-dependent information is already encoded in the neuronal activity. For example, if the neuronal tuning curve and the astrocytic-induced change in neural function are similarly modulated by the animal’s position, no additional dependence of neuronal function by position would be introduced by astrocyte-neuron interactions. Conversely, our findings imply that astrocytic calcium dynamics carrying *complementary information* to that of neurons enable modulations of synaptic transmission and neuronal gain which could increase the computational capability of neural circuits ^41, 42^. For example, changing the gain of neurons with a coordinate system complementary to that regulating its tuning function has been shown to endow neural networks with richer computations ^42, 43^. Moreover, targeted dynamic control of neural excitability (e.g., changing the gain of a subset of neurons in the network rather than the whole network) can greatly increase the dynamic repertoire and coding capabilities of circuits, for example by making it possible to reach different attractors from a similar set of initial conditions^44^. We thus propose that the complementary place-dependence of the astrocytic calcium dynamics and the place-dependence of astrocytic-neuron interactions reported here facilitate the emergence of dynamic, context-dependent changes in population coding of CA1 neurons. Our work calls for a re-examination of the theory of place coding and of brain population codes in light of the opportunities offered by complementary astrocytic information coding. We propose that the complementary regulation of astrocytic calcium activity and of its interaction with neurons may reflect a general principle of how the brain encodes information. This conclusion may extend beyond the hippocampus and spatial navigation to other brain regions and cognitive tasks and it will need to be included in the conceptualization of brain function.

## Methods

### Animals

All experiments involving animals were approved by the National Council on Animal Care of the Italian Ministry of Health and carried out in accordance with the guidelines established by the European Communities Council Directive (authorization 61/2019-PR). From postnatal day 30, animals were separated from the original cage and housed in groups of up to five littermates per cage with *ad libitum* access to food and water in a 12-hour light-dark cycle. Experimental procedures were conducted on animals older than 10 weeks. The number of animals used for each experimental data set is specified in the text or in the figure legends.

### AAV injection and chronic hippocampal window surgery

Astrocytic-specific GCaMP6f expression was obtained using pZac2.1 gfaABC1D-cyto-GCaMP6f (Addgene viral prep # 52925-AAV5 a gift from Dr. Khakh, ^4, 20^). Neuronal-specific jRCaMP1a expression was obtained using pAAV-CAMKII-jRCaMP1a (kindly provided by Dr. O. Yizhar) which was then packaged as AAV serotype 1-2 viral particles ^45^.

Male C57Bl6/j mice were anesthetized with 2% isoflurane/0.8% oxygen, placed into a stereotaxic apparatus (Stoelting Co, Wood Dale, IL), and maintained on a warm platform at 37 °C for the whole duration of the anesthesia. Before surgery, a bolus of Dexamethasone (4 mg/kg, Dexadreson, MSD Animal Health, Milan, Italy) was provided with an intramuscular injection. After scalp incision, a 0.5 mm craniotomy was drilled on the right hemisphere (1.75 mm posterior, 1.35 mm lateral to bregma) and the AAV-loaded micropipette was lowered into the hippocampal CA1 region (1.40 mm deep to bregma). 800 nL of AAV solution was injected at 100 nL/min by means of a hydraulic injection apparatus driven by a syringe pump (UltraMicroPump, WPI, Sarasota, FL). Following the viral injection, a stainless-steel screw was implanted on the cranium of the left hemisphere and a chronic hippocampal window was implanted similarly to ^17, 46^. A drill was used to open a 3 mm craniotomy centered at coordinates 2.00 mm posterior and 1.80 mm lateral to bregma. The dura was removed using fine forceps, and the cortical tissue overlaying the hippocampus was carefully aspirated using a blunt needle coupled to a vacuum pump. During aspiration, the exposed tissue was continuously irrigated with HEPES-buffered artificial cerebrospinal fluid (ACSF). Aspiration was stopped once the thin fibers of the external capsule were visible. A cylindrical cannula-based optical window was fitted to the craniotomy in contact to the external capsule and a thin layer of silicone elastomer (Kwik-Sil, World Precision Instruments, Sarasota, FL) was used to surround the interface between the brain tissue and the steel surface of the optical window. A custom stainless-steel headplate was attached to the skull using epoxy glue. All the components were secured in place using black dental cement and the scalp incision was sutured to adhere to the implant. Animals received an intraperitoneal bolus of antibiotic (BAYTRIL, Bayer, Germany) at the end of the surgery.

Optical windows were composed of a thin-walled stainless-steel cannula segment (OD, 3 mm; ID, 2.77 mm; height, 1.50 - 1.60 mm). A 3.00 mm diameter round coverslip was attached to one end of the cannula using UV curable optical epoxy (Norland optical adhesive 63, Norland, Cranbury, NJ). Sharp edges and bonding residues were smoothed using a diamond-coated cutter.

### Virtual reality

A custom virtual reality setup was implemented using the open-source 3D creation suite Blender (blender.org, version 2.78c). Virtual environment rendering was performed using the Blender Game Engine and displayed at video rate (60 Hz). The virtual environment was a linear corridor with the proximal walls characterized by three different white textures (vertical lines, mesh, and circles) on a black background. Distal walls were colored in green and labeled with a black cross. The corridor was 180 cm long and 9 cm wide. The character avatar was a sphere of radius 2 cm with a rectangular cuboid protruding at the equator parallel to the corridor floor (cuboid dimension: x = 5 cm, y = 1 cm, z = 1 cm). The cuboid acted as a virtual touch sensor with the environment. The character point of view (220° horizontal, 80° vertical) was rendered through a composite tiling of five thin bezel-led screens. The virtual corridor implementation described above was used for both monodirectional and bidirectional navigation. In monodirectional virtual navigation, mice navigated the environment running on a custom 3D printed wheel (radius 8 cm, width 9 cm). An optical rotary encoder (Avago AEDB-9140-A14, Broadcom Inc., San Jose, CA) captured motion and a single board microcontroller (Arduino Uno R3, Arduino, Ivrea, Italy) performed USB-HID-compliant conversion to a serial mouse input. In bidirectional virtual navigation, mice navigated the environment using an air-suspended Styrofoam ball (radius, 10 cm) and a Bluetooth optical mouse (M170, Logitech, Lausanne, Switzerland) was used to read the vertical and horizontal displacement. In both monodirectional and bidirectional navigation, physical motion of the input devices was mapped 1:1 to the virtual environment. To motivate corridor navigation, mice received ∼ 4 µl water rewards upon reaching specific locations. Rewards were delivered through a custom steel lick-port controlled by a solenoid valve (00431960, Christian Bürkert GmbH & Co., Ingelfingen, Germany) and licks were monitored using a capacitive sensor (MTCH102, Microchip Technology Inc., Chandler, AZ). In monodirectional virtual navigation, rewards were delivered at 115 cm and the mouse was teleported to the beginning of the corridor after reaching the end of the track (inter trial timeout interval 5 s). If the mouse didn’t reach the end of the corridor within 120 s, the trial was automatically terminated and the mouse was teleported to the beginning of the corridor after an inter-trial timeout. For bidirectional navigation, rewards were delivered at opposite ends of the track. After getting a reward, the mouse had to reach the opposite end of the virtual corridor to receive the next reward. Virtual reality rendering and two-photon imaging acquisition ran on asynchronous clocks while the command signal of the slow galvanometer was used to synchronize the imaging acquisitions with behavior.

### Two-photon imaging during virtual navigation

Two-photon calcium imaging was performed using an Ultima Investigator or an Ultima II scanheads (Bruker Corporation, Milan, Italy) equipped with raster scanning galvanometers (6 mm or 3 mm), a 16x/0.8 NA objective (Nikon, Milan, Italy), and multi-alkali photomultiplier tubes. For GCaMP6f imaging, the excitation source was a Chameleon Ultra pulsed laser tuned at 920 nm (80 MHz repetition rate, Coherent, Milan, Italy). Simultaneous GCaMP6f and jRCaMP1a imaging was performed with two optical path configurations. On the Ultima Investigator, two pulsed laser sources were combined through a dichroic mirror (zt98rdc-UF1, Chroma Technology Co., Bellow Falls, VT; λ_1_= 920 nm, Alcor 920 fiber laser - 80 MHz repetition rate, Spark Lasers, Martillac, France; λ_2_= 1060 nm, Chameleon Ultra II - 80 MHz repetition rate, Coherent, Milan, Italy). On the Ultima II, two orthogonally polarized pulsed laser sources were combined through a polarizing beam splitter (05FC16PB.5, Newport; λ_1_= 920 nm, Chameleon Ultra II; λ_2_= 1100 nm, Chameleon Discovery - 80 MHz repetition rate, Coherent, Milan, Italy). Laser beam intensity was adjusted using Pockel cells (Conoptics Inc, Danbury, USA). Imaging average power at the objective outlet was ∼ 80 - 110 mW. Fluorescence emission was collected by multi-alkali PMT detectors downstream of appropriate emission filters (525/70 nm for GCaMP6f, 595/50 nm for jRCaMP1a). Detector signals were digitalized at 12 bits. Imaging sessions were conducted in raster scanning mode at ∼ 3 Hz using 5x optical zooming factor. Images contained 256 pixels x 256 pixels field of view (pixel dwell-time, 4 µ s; Investigator: pixel size, 0.63 µm; Ultima II: pixel size, 0.51 µm).

One or two weeks after surgery the animals were set on a water restricted schedule, receiving approximately 1 ml of water per day. Weight was monitored daily, and remained between 80 – 90 % of the starting weight throughout all procedures. Mouse habituation to the experimenter (handling) started two days after water scheduling and lasted for a minimum of two sessions. Following handling, mice were habituated to the virtual reality setup in successive training sessions. Starting from the second habituation session, the animals were head-tethered for a progressively increasing amount of time, reaching 1 hour in approximately one week. During virtual reality training sessions, mice were exposed to the noise generated by the two-photon imaging setup (e.g., galvanometer scanning noise, shutter noise). Training in the virtual environment lasted until animals routinely ran along the linear track. On experimental days, mice were head-tethered, and the virtual reality session started after a suitable field of view was identified. At the end of each imaging session, animals were returned to their home cage.

### Data analysis

#### Motion correction, image segmentation, and trace extraction

Analysis was performed using Python 3.6 (python.org) and custom code. t-series were pre-processed to correct motion artifacts using an open-source implementation of up-sampled phase cross-correlation ^47, 48^. Regions (typically at the edges of the field of view) within which artifacts could not be corrected were not considered for analysis.

For astrocytic recordings, ROI segmentation was performed on median projections after motion correction using manual annotation. Astrocytic ROIs were classified as soma or process according to visible anatomic features. For each ROI, fluorescence signals were computed as:

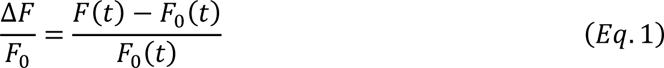

where F(t) is the average fluorescence signal of a given ROI at time t, and F_0_(t) is the baseline fluorescence, computed as the 20^th^ percentile of the average fluorescence intensity in a 30 s-long rolling window centered in t.

For neuronal recordings, cell identification was performed on the median temporal projection of each t-series, after motion correction, by identifying rectangular boxes containing the neuronal soma of the identified neuron, as in ^49^. Only pixels with signal-to-noise (SNR) value greater than the 80^th^ percentile of the SNR distribution were considered as part of the ROI corresponding to the considered rectangular box. The neuropil signal was computed as the average trace of all pixels in the time series not belonging to bounding boxes. This value was multiplied by a factor r = 0.7 ^18^ and then subtracted from each fluorescence trace. ΔF/F_0_ traces were computed as:

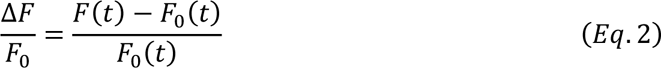

where F(t) is the neuropil-subtracted fluorescence trace signal at time t, and F_0_(t) is the baseline trace computed as 20^th^ percentile of the average intensities in a 10 s rolling window centered in t.

#### Identification of calcium events

For both astrocytic and neuronal fluorescence traces, extraction of statistically significant calcium events was performed on ΔF/F_0_ traces via modified implementation of the algorithm described in^50^. For all subsequent analysis, an event trace was obtained from the ΔF/F_0_ trace by setting all fluorescence values outside of those belonging to positive events to 0.

#### Identification of reliable spatial modulation of calcium signals

To evaluate if and how position in the virtual corridor modulated calcium signals, we applied two basic requirements: that activity carried significant information about position, and that the spatial modulation properties were reliably reproducible across subsets of trials. We restricted the analysis to running-trials, defined as consecutive frames of forward locomotion in which mouse speed was greater than 1 cm/s. Calcium responses were considered with reliable spatial information if they matched both of the following criteria: *i)* response field reliability was greater than 0 (see *Spatial reliability of calcium responses*); and *ii)* mutual information between position and calcium event trace was significant (see *Spatial information in calcium signals*). The same criteria were applied to astrocytic ROIs and neuronal ROIs.

#### Analysis of calcium responses during virtual navigation

Analysis was performed on all running-trials, binning the length of the virtual corridor (number of spatial bins, 80; bin width, 2.25 cm). For each ROI, the occupancy map was built by computing the total amount of time spent in each spatial bin. The activity map was computed as the average fluorescence value in each spatial bin. Both the activity map and the occupancy map were normalized to sum 1 and convolved with a Gaussian kernel (width of the Gaussian, σ, was equal to 3 spatial bins, which corresponded to 6.75 cm). The response profile of an ROI, *RP*, was defined as the ratio of the activity map over the occupancy map for that ROI. For each *RP*, we identified a response field, RF, as follows: *i)* the array of local maxima greater than the 25^th^ percentile of the response profile values was selected, C = (c0, c1, …, cn) ; *ii)* the elements of *C* were used to initialize the fitting of the sum of a set of n parametrized Gaussian functions, with mean at one of the elements of C, amplitude (*a*) at 0 ≤ a ≤ 1, and standard deviation (σ) at 0 ≤ σ ≤ 90 cm; *iii)* this set of Gaussian functions was fitted to the response profile to solve a non-linear least squares problem (curve_fit function from ^51^); and *iv)* the response field was defined as the Gaussian with the highest amplitude and the response field width was defined as 2σi.

Thus:

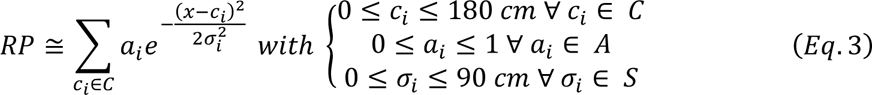

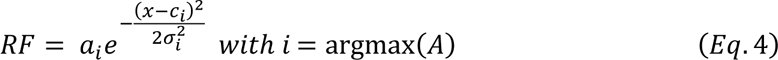

#### Spatial reliability of calcium responses

To quantify the spatial reliability of response fields, we computed response profiles subsampling either odd or even running-trials. For either fraction of running-trials we estimated response field center (c_odd_, c_even_) and response field half-width (σ_odd_, σ_even_). We quantified spatial reliability of calcium responses as a similarity index, where the absolute difference of response field centers, obtained with either fractions of the running-trials, was inversely weighted by the most conservative estimate of response field width:

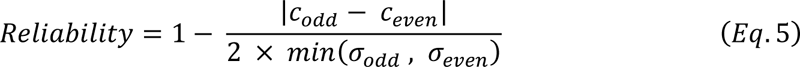

ROIs with reliability greater than 0 were considered reliable (Extended Data Fig. 1).

#### Spatial information in calcium signals

We used information theory to quantify our information gain (or reduction of uncertainty) about position obtained by knowing the calcium response ^22, 52^. We computed the mutual information, *I(S;R*), between position in the linear track, stimulus (S), and the calcium event trace, response (R), as follows:

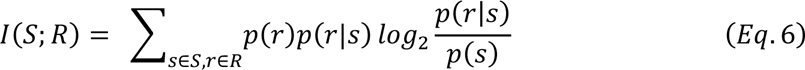

with S and *R* representing the arrays of all possible discrete values of stimulus or response, *p(s)* the probability of the stimulus *s*, *p(r)* the probability of the response r across all trials to any stimulus, and *p(r|s)* the conditional probability of the responses *r* given presentation of stimulus *s*. We characterized the effects of discretization on the estimates of mutual information, computing mutual information while changing the number of discrete states (N) for both *S* (N_S_ = 4, 8, 12, 16, 20, 24, 40, 60, 80, 100, 160) and *R* (N_R_ = 2, 3, 4, 5, 8, 10, 20). Statistical significance of mutual information was tested using a non-parametric permutation test. We randomly permuted the calcium event trace 10^4^ times, removing any relationship between *R* and *S*. We used shuffled traces to compute a null distribution of mutual information values. A mutual information value was considered significant if greater than the 95^th^ percentile of the null distribution. Mutual information values were conservatively corrected for limited-sampling bias subtracting the mean value of the null distribution ^53, 54^. The results of this analysis for astrocytic ROIs are reported in Extended Data Fig. 1. To allow robust estimates of mutual information values while preserving adequately fine discretization of position, we used N_s_ = 12 throughout the manuscript. For single ROIs analysis reported in figures Fig. 1, Fig. 2, and Fig. 4, we used N_R_ = 4 to discretize astrocytic calcium event traces and N_R_ = 2 for binarized neuronal event traces (setting to 1 all the non-zero values as in ^55^).

#### Directionality of astrocytic spatial responses

In experiments where the mouse performed bidirectional navigation, astrocytic ROIs could be spatially-modulated in either running direction. To quantify whether responses were direction selective, we computed the directionality index (DI) as:

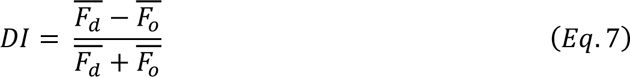

where *F̄*_*d*_ was the average of Δ*F*/*F_0_* inside the response field, and *^F̅^_o_* was the average of Δ*F*/*F_0_* at the same response field while running in the opposite direction. *DI* > 0 indicated that average response at the response field was direction-selective. We compared the distribution of *DI* values for all spatially-modulated ROIs with surrogate data. To this end, we randomly selected one of the informative ROIs and computed *DI* after applying a random shift of response field position along the linear track while preserving its width. We repeated this operation 10^5^ times, obtaining a distribution of *DI* values representing the occurrence of *DI* values at any spatial location as wide as a response field.

#### Population analysis using Mutual Information

For experiments in which we simultaneously recorded astrocytic and neuronal calcium activity, we used all running-trials to compute the mutual information about animals’ position obtained by observing the calcium signals of a pair of simultaneously recorded ROIs. Results are reported as a function of pair composition, with pairs containing either two astrocytic ROIs, two neuronal ROIs, or one element of each type.

Mutual information between the spatial position, *S*, and the array of joint responses for a pair of ROIs, *R* = (*R_1_*,*R_2_*), was computed as ^27^:

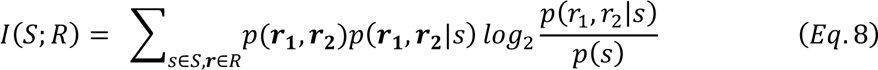

where *p(s)* is the probability stimulus *s*, *p(r_1_,r_2_)* is the probability of joint responses *r_1_* and *r_2_* across all trials to any stimulus, and *p(r_1_,r_2_|s)* is the conditional probability of the joint responses *r_1_* and *r_2_* given presentation of stimulus *s*.

For consistency with single-ROI analysis, spatial position was discretized with N_s_ = 12. To allow consistent scaling of probability spaces and comparable information values, the astrocytic calcium event trace was binned with N_R_ = 2 (we verified that the main conclusions were maintained when using N_R_ = 3 and N_R_ = 4), and N_R_ = 2 for neuronal calcium event trace discretization, as described for single neuron analysis. We corrected for the limited sampling bias as described in refs ^26, 58^.

To quantify whether the within-trial correlations of a given ROI pair enhanced the amount of position information carried by the pair, we used trial-shuffling to disrupt the within-trial correlations between ROIs while keeping intact the spatial position information of individual ROIs. Within subsets of trials with the same position bin, we generated pseudo-population responses by independently combining shuffled identities of trials for each ROI. Thus, responses of individual ROIs to the spatial position were maintained while within-trial correlations between ROIs were disrupted. We computed 100 trial-shuffling estimates of mutual information, I(S;R)_trial-shuffled_, for calcium responses at fixed position. A pair was classified as having information enhanced by correlations, if I(S;R) was greater than the 95^th^ percentile of the corresponding I(S;R)_trial-shuffled_ distribution.

#### Information Breakdown

We performed information breakdown analysis ^26, 27^. We decomposed spatial information carried by a pair of ROIs, I(S;R), into 4 terms. Each term expressed a different contribution carried by correlations to the information between the ROIs. The decomposition is as follows:

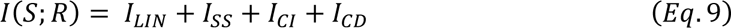

*I_LIN_*, the mutual information linear term, is the sum of the information provided by each ROI. *I_SS_* (signal similarity term) is a non-positive term quantifying the decrease of information (amount of redundancy) due to signals correlation caused by correlations between the trial-averaged spatial position profiles of the calcium signals of the two ROIs. *I_CI_* (stimulus independent correlation) is a term that can be either positive, null, or negative and that quantifies the contribution of stimulus-independent correlations. *I_CI_* is negative if noise and signal correlations have the same signs and positive otherwise. *I_CD_* (stimulus-dependent correlational term) is a non-negative term that quantifies the amount of information, above and beyond that carried by the responses of individual ROIs carried by stimulus modulation of noise correlation strength. Although *I_CD_* is strictly non-negative, *I_CD_* values could occasionally become slightly negative due to quadratic extrapolation bias correction.

The above calculations of I(S;R) were conducted with a bias correction procedure that, with the typical number of trials per spatial location represented in our data, was shown to be accurate for removing the limited sampling bias ^59^.

#### Position-dependent correlation

To measure whether correlation between pairs of neuronal and astrocytic ROIs was position-dependent, we computed pairwise Pearson’s correlations between calcium signals sampled inside and outside the response fields. On average, response fields were smaller than half the linear track, thus either set of observations, inside or outside the response field, could contain uneven amounts of datapoints. To compensate for the unbalanced numerosity, we resampled the same number of points found in the smaller set, while preserving temporal ordering. We then computed Pearson’s correlation between the two vectors. For each pair of ROIs, we computed the average Pearson’s correlation with 100 iterations of this procedure. We repeated this procedure inside both astrocytic fields and neuronal response fields.

#### Population analysis using SVM decoder of spatial position

To decode animals’ position from a population of ROIs, we trained an SVM classifier ^60–62^. We performed decoding analysis on three datasets: i) astrocytic signals during monodirectional virtual navigation; ii) astrocytic signals during bidirectional virtual navigation; iii) simultaneous recording of astrocytic and neuronal signals during monodirectional virtual navigation. Experimental sessions were considered independently. We evaluated decoding performance as a function of decoding granularity, *G*, i.e., the number of spatial bins we used to discretize the linear track. For monodirectional virtual navigation, we used G = (4, 8, 12, 16, 20, 24), and for bidirectional virtual navigation, for which there was a limited number of running trials, we used G = (4, 8, 12, 16).

We used experimental session as the n-dimensional array of calcium event traces (n = number of ROIs) to decode discretized positions along the virtual linear track at each time point. Each experimental session was composed of a set of T_exp_ observations (X_i_,y_i_), where X_i_ is the n-dimensional array of the calcium activity of the n ROIs, whereas y_i_ corresponds to the discretized spatial position. For each granularity, we trained and tested the SVM using 10-fold cross-validation procedure on each experimental session independently. Predictions of the decoder for each of the 10-folds used as test were then collected to compute the overall performance of the decoder.

For each granularity, we measured decoding performance computing decoded information, as the mutual information between predicted and real spatial position ^22^:

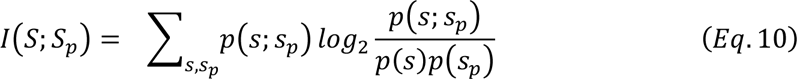

where *s_p_* denotes the decoded spatial position (with the SVM method described above) from the population response vector in each trial, *s* is the actual spatial position of the animal, and p(s; s_p_) is the decoder’s confusion matrix obtained from the predictions of the 10-folds cross-validation test-set. We corrected mutual information measures for the limited sampling bias as described in refs ^53, 54, 59^.

Decoding performance was also computed as decoding accuracy (fraction of correct predictions):

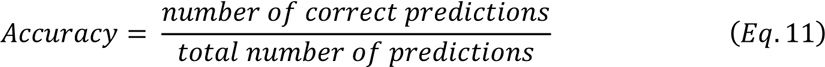

To assess the statistical significance of decoding results, we trained and tested the decoder on each experimental session after randomly permuting position and responses. This procedure removed all information about position carried in the responses. We performed 10^3^ random permutations for each granularity and population type. We then used the distribution of information values on permuted data as the null hypothesis distribution for the one-tailed non-parametric permutation test of whether information was significantly larger than zero. We repeated this procedure separately for each granularity.

To assess if the correlations among neurons and/or astrocytes increased the amount of spatial information, we disrupted across-neuron correlations by randomly shuffling, separately for each ROI, the order of trials with the same position category. We performed 500-trial shuffling for each granularity and population type. We then used the trial-shuffled distribution as the null hypothesis distribution for the one-tailed non-parametric permutation test of whether the information in the real population vector (which includes correlations) is significantly higher than that obtained when correlations are removed.

#### Decoding error analysis

We investigated classification errors made by the decoder for each decoding granularity. We considered only misclassified samples in the test set and we measured the distance between the position predicted by the decoder and the ground truth position. We computed the frequency histogram of these deviations from the ground truth, and fitted a Gaussian curve ^51^ using non-linear least squares. For each histogram, we computed R^2^ score to quantify the fitting performance.

#### Statistics

Significance threshold for statistical testing was always set at 0.05. No statistical methods were used to pre-determine sample size, but sample size was chosen based on previous studies ^17, 21, 46^. Statistical analysis was performed using Python (SciPy 0.24, NumPy 1.19, statsmodels 0.9), or the InfoToolbox library ^26^ available for Matlab (MathWorks R2019b). A Python 3 ^63^ (version 3.6) front-end was used for execution. To test for normality, either a Shapiro-Wilks (for N ≤ 30) or a D’Agostino K-squared test (for N > 30) was run on each experimental sample. When comparing two paired populations of data, a paired t-test or Wilcoxon signed-rank test were used to calculate statistical significance (for normal and non-normal distributions, respectively). Independent samples t-test and two-sample Kolmogorov-Smirnov test or Wilcoxon rank-sum test were used for unpaired comparisons of normally and non-normally distributed data, respectively. Bonferroni correction was applied to correct for the multiple testing problem when appropriate. Surrogate data testing was performed as described in the specific methods sections. All tests were two-sided, unless otherwise stated. When reporting descriptive statistics of data distributions, we used either the mean ± standard deviation (mean ± s.d.) for normal data or the median ± median absolute deviation (median ± m.a.d.) for non-normal data. Datasets reporting average values across experimental sessions were presented as mean ± standard error of the mean (mean ± s.e.m.).

### Data availability

The data are available from the corresponding author upon request.

### Code availability

The code is available from the corresponding author upon request, and it will be shared upon publication.

## Acknowledgments

We thank C. Harvey, G. Edgerton, B. Mensh for helpful comments on the manuscript, L. Sità for technical help, O. Yizhar for sharing the jRCaMP1a construct, A. Attardo for help with surgical procedures, and A. Contestabile for help with the virus production.

## Author contributions

SC built the experimental set up and performed experiments and analyses. JB performed analyses. SR contributed to two-color imaging experiments. SP developed and supervised analyses. TF conceived and supervised the project. TF, SC, and SP wrote the paper with contribution from all authors.

## Extended data

This work has extended data in Figures 1 to 13 and Tables 1 to 6.

## Extended data

**Extended data Figure 1.**
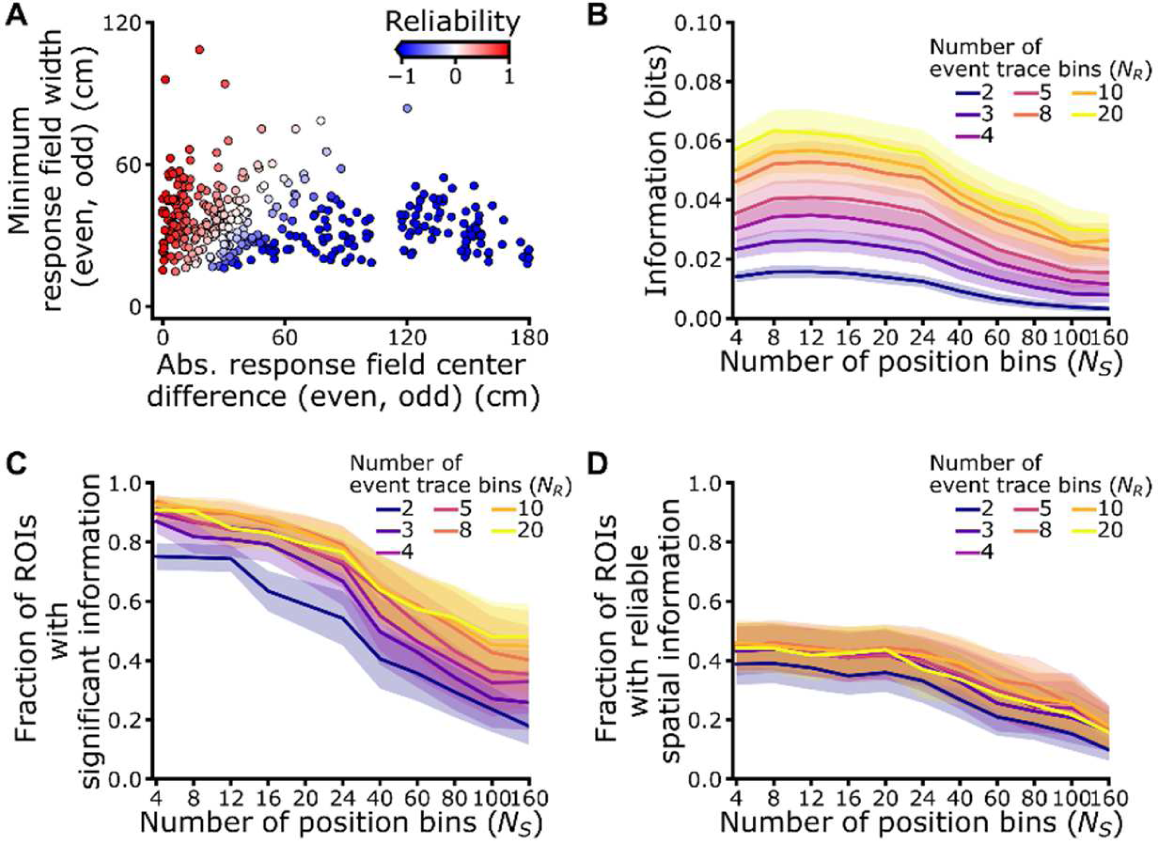
Identification of reliable spatial modulation of astrocytic calcium signals. (A) Minimum response field width between even and odd trials as a function of the difference in place field position. The pseudocolor scale indicates reliability of the response (see Methods). (B-C) Mutual information values (B) and fraction of ROIs showing significant spatial information (C) as a function of the number of bins for the stimulus (animals’ position in the linear track). Colors indicate different binning of the response (calcium event trace). Mutual information values were bias-corrected using bootstrap method (10^4^ iterations). Significance level for information content was set at p < 0.05. (D) Fraction of ROIs with reliable spatial information as a function of the number of bins for the stimulus. Colors indicate different binning of the response. Data in (B-D) are presented as mean ± s.e.m. from 7 imaging sessions in 3 animals.

**Extended data Figure 2.**
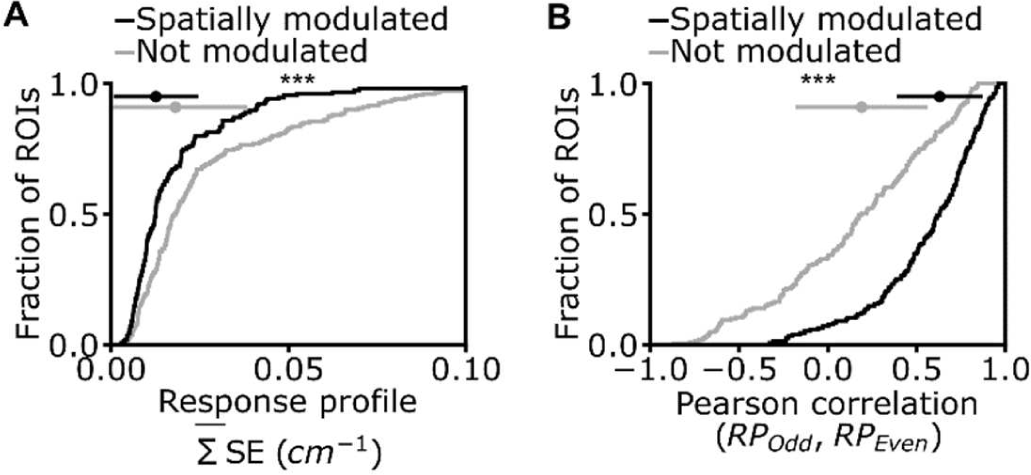
Reliable spatial modulation of astrocytic calcium signals. (A) Cumulative distribution of the mean standard error (SE) of the response profile in astrocytic ROIs (median ± m.a.d. 1.3E-2 ± 1.2E-2 cm^-1^, N = 155 out of 356 total ROIs, for ROIs with reliable spatial information, black; 1.8E-2 ± 2.0E-2 cm^-1^, N = 201 out of 356 total ROIs, for not-modulated ROIs, grey: p = 1E-5, Kolmogorov-Smirnov test). (B) Cumulative distribution of Pearson correlation values between astrocytic response profiles in even and odd trials (median ± m.a.d. 0.63 ± 0.24, N = 155 out of 356 total ROIs for ROIs with reliable spatial information, black; 0.19 ± 0.37, N = 201 out of 356 total ROIs, for not-modulated ROIs, grey; p = 5E-14, Kolmogorov-Smirnov test). In all panels, data from 7 imaging sessions in 3 animals.

**Extended data Figure 3.**
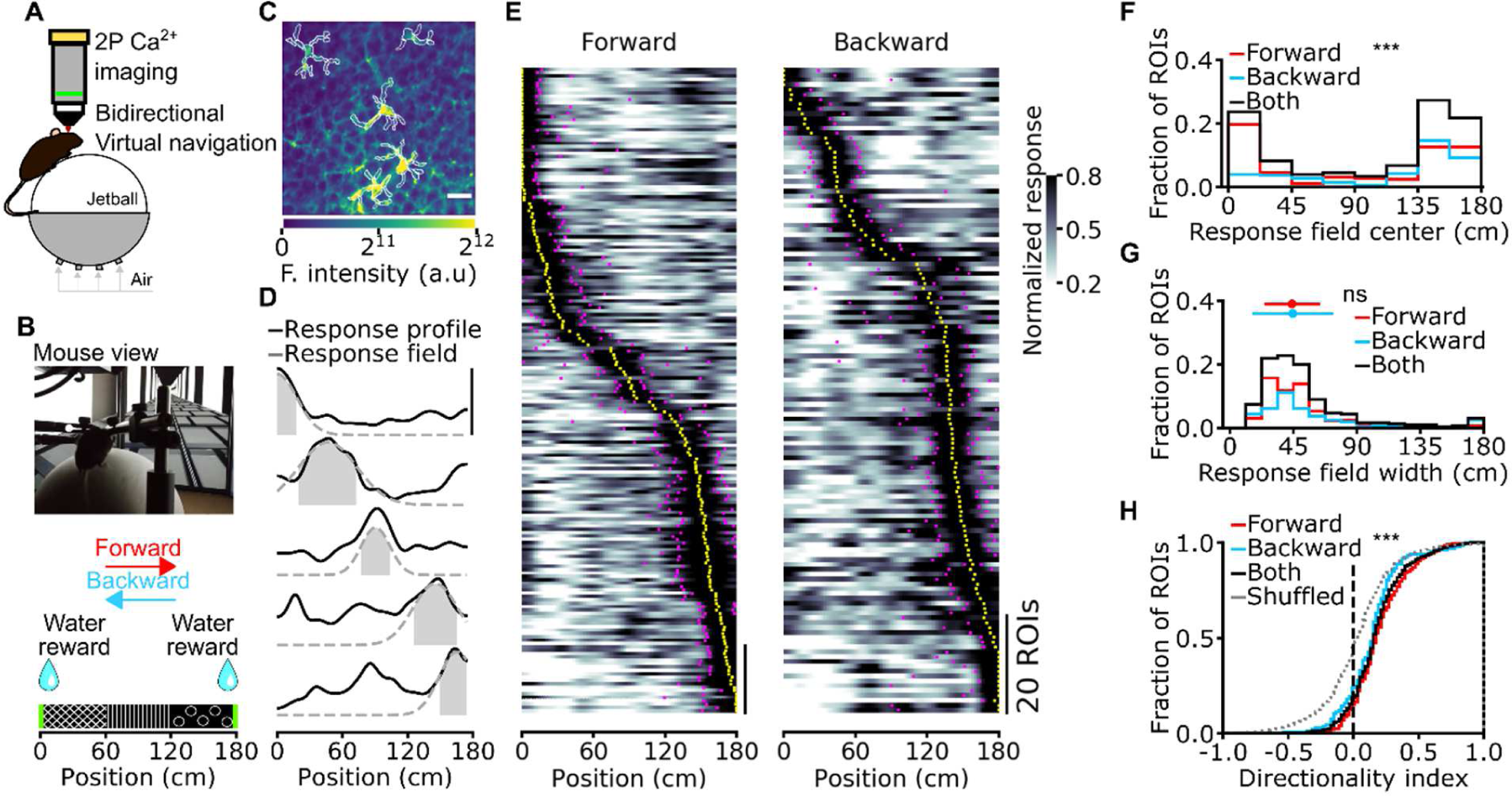
Calcium signals of CA1 astrocytes encode direction-selective spatial information during virtual bidirectional navigation. (A) Two-photon functional imaging of CA1 astrocytes is performed during bidirectional virtual navigation. (B) Head-restrained mice run on an air-suspended spherical treadmill in a linear virtual track in both forward and backward directions. Water rewards are delivered at either end of the virtual corridor. (C) Median projection of GCaMP6f-labeled astrocytes in the CA1 pyramidal layer. White lines indicate segmented ROIs (scale: 20 µ m). (D) Calcium signals for five representative astrocytic ROIs reliably encoding spatial information across the corridor length. Solid black lines indicate the average astrocytic calcium response across trials as a function of spatial position. Dashed grey lines and filled grey areas indicate the Gaussian fitting function and the response field width (see Methods), respectively. (E) Normalized astrocytic calcium responses as a function of position for astrocytic ROIs with reliable spatial information. Trials are divided according to running direction (forward and backward). For forward trials, informative ROIs are N = 192 out of 648 total ROIs, mean ± s.d.: 29 ± 13 %; for backward trials, informative ROIs are N = 133 out of 648 ROIs, mean ± s.d.: 20 ± 13 %, p = 0.09, Wilcoxon signed rank test. Scale bar: 20 ROIs. Yellow dots indicate the center position of the response field, the magenta dots indicate the width of the field response. (F) Distributions of astrocytic response field position for forward and backward running direction. Median ± m.a.d. 93 ± 66 cm, N = 192 out of 648 total ROIs for the forward direction; 138 ± 47 cm N = 133 out of 648 total ROIs for the backward direction; p = 9E-7, Kolmogorov-Smirnov test). (G) Distributions of response field width for the forward and backward running direction (response field width, 44 ± 19 cm, N = 192 out of 648 total ROIs for the for forward direction; response field width, 44 ± 28 cm, N = 133 out of 648 total ROIs for the backward direction; p = 0.34, Wilcoxon rank sums test). (H) Directionality index for forward and backward running directions (directionality index, 0.18 ± 0.16, N = 192 out of 648 total ROIs for forward trials; directionality index, 0.16 ± 0.16, N = 133 out of 648 total ROIs for backward trials; p = 8E-19 and p = 2E-8, respectively, Kolmogorov-Smirnov test *vs* shuffled distribution). In all panels, data from 18 imaging sessions in 4 animals.

**Extended data Figure 4.**
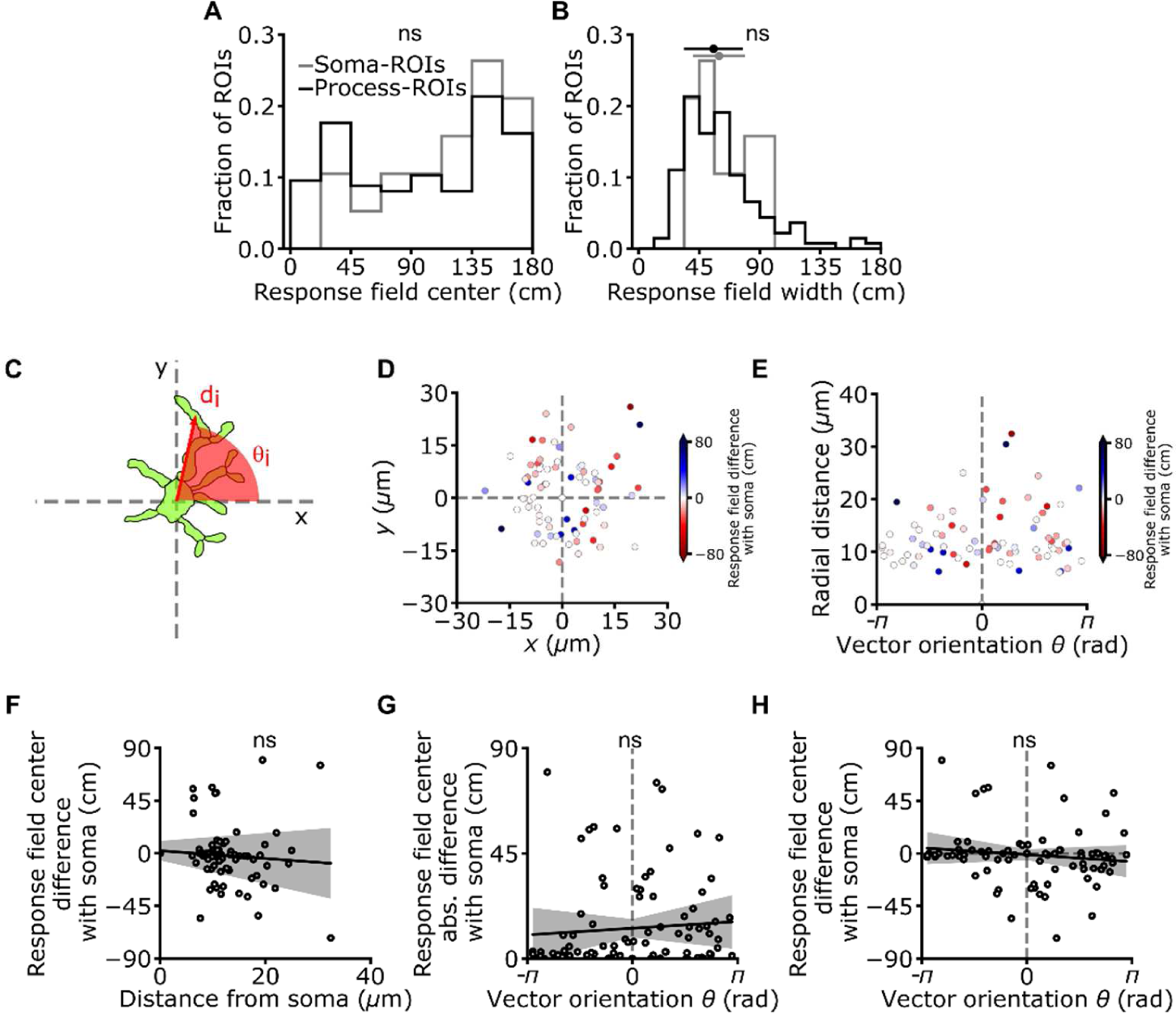
Anatomical organization of subcellularly localized astrocytic calcium signals. (A) Distribution of field position for soma-ROIs and process-ROIs (p = 0.36, Kolmogorov-Smirnov test). (B) Distribution of response field width for astrocytic soma-ROIs and process-ROIs (median width for soma-ROIs: 60 ± 19 cm; median width for process-ROIs: 56 ± 22 cm, p = 0.36, Wilcoxon Rank sums test). (C) For each pair of ROIs within a given astrocyte, the distance (d) between the centers of two ROIs and the angle between the line connecting the two ROI centers and the X axis are calculated. Only astrocytes showing significant spatial modulation in the soma and at least one process were used for this analysis. (D, E) Difference in field position of a process with respect to the field position of its corresponding soma, expressed as function of Cartesian (D) and polar (E) coordinates of the ROI centers. (F) Difference in response field position of a process with respect to the field position of its corresponding soma as a function of the process distance from cell soma (R^2^ = 0.01, p = 3.3E-1, Wald test, data from 19 cells from 7 imaging sessions on 3 animals). (G, H) Absolute value (G) or signed (H) difference in response field position of a process-ROI with respect to the field position of its corresponding soma as a function of the process angular coordinate (absolute value of difference in response field R^2^ = 0.01, p = 4.8E-1, Wald test; signed value of difference in response field R^2^ = 0.01, p = 4.1E-1, Wald test, data from 19 cells from 7 imaging sessions on 3 animals).

**Extended data Figure 5.**
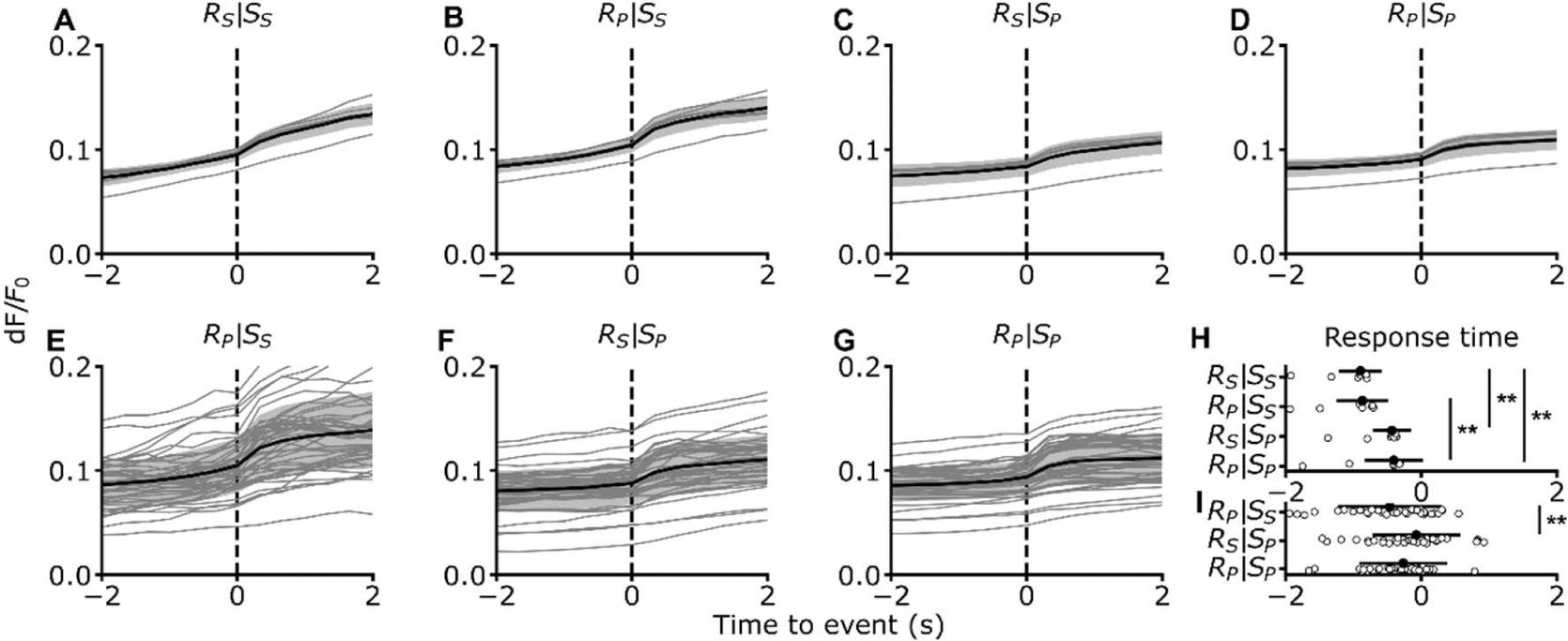
Temporal relationships among subcellularly localized astrocytic calcium signals. (A) Event triggered average of astrocytic calcium responses. Calcium responses of putative receiver (R) ROIs are aligned to calcium events of putative source (S) ROIs according to anatomic identities of ROIs (e.g., somatic receiver ROIs and somatic source ROIs). Data from 7 imaging sessions in 3 animals. Black line indicates the mean, shaded area the standard deviation. (B-D) Same as in (A) for pairs of process receiver and somatic source (B), somatic receiver and process source (C), and process receiver and process source (D). (E-G) Same as in (B-D) but for pairs of ROIs belonging to the same astrocyte (N = 46 astrocytes from 7 imaging sessions in 3 animals). (H) Response time (see Methods) for signals shown in (A-D). p = 6E-4, Friedman test with Nemenyi post-hoc correction. (I) Response time for signals shown in (E-G). p = 7E-3, Friedman test with Nemenyi post-hoc correction.

**Extended data Figure 6.**
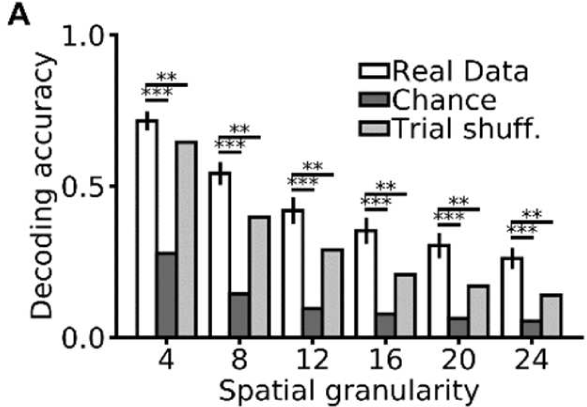
Decoding animal’s position from astrocytic calcium signals in the unidirectional virtual navigation task. (A) Decoding accuracy as a function of spatial granularity on real (white), chance (dark gray), and trial-shuffled (grey) data (see Methods). Data are presented as mean ± s.e.m. from 7 imaging sessions on 3 animals, see also Extended Data Table 2.

**Extended data Figure 7.**
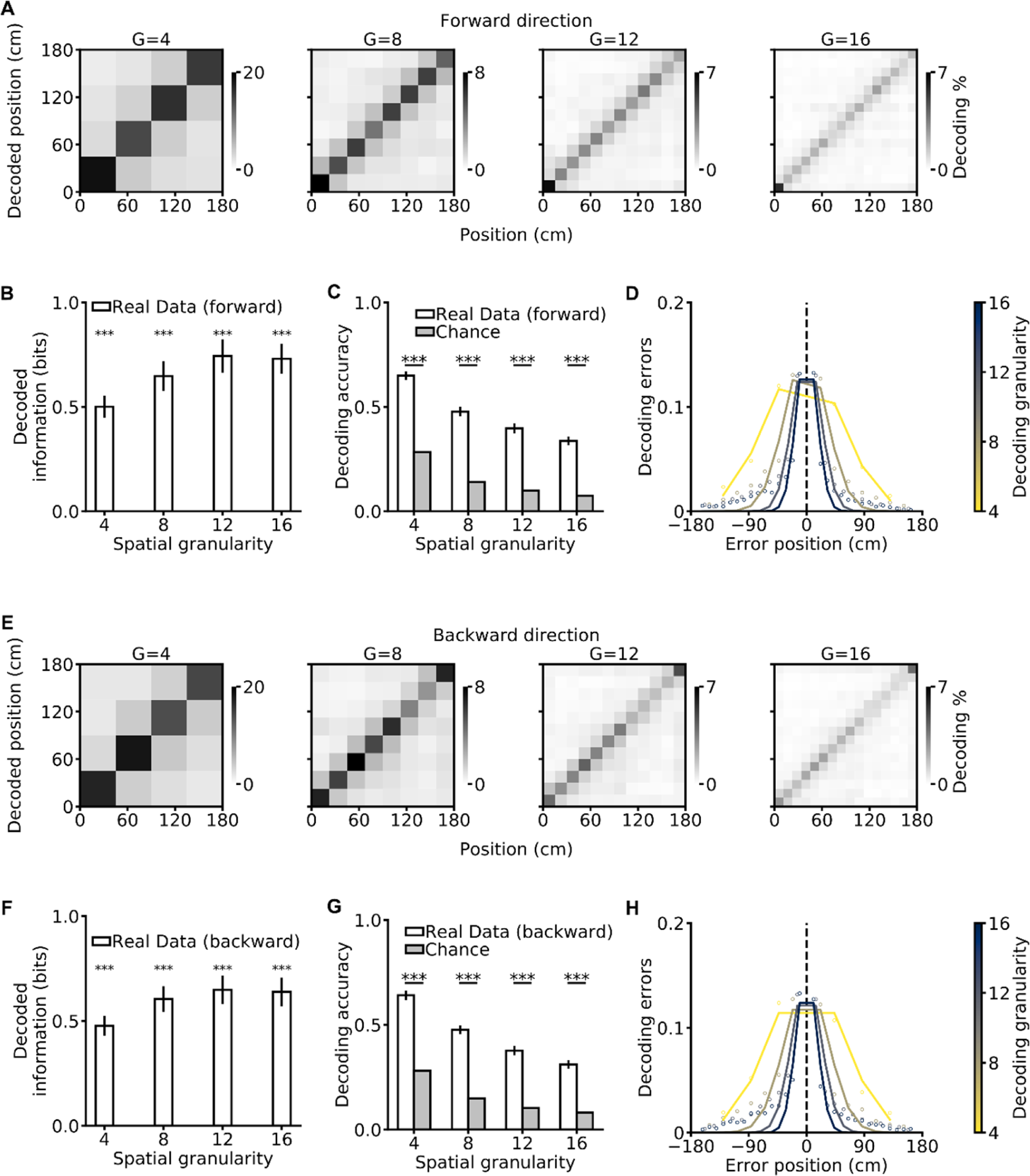
Decoding animal’s position from astrocytic calcium signals in the bidirectional virtual navigation task. (A) Confusion matrices of a SVM classifier for different spatial granularities (G = 4, 8, 12, 16) for trials in which the mouse was running in the forward direction (forward). The actual position of the animal is shown on the x-axis, the decoded position on the y-axis. Grey scale indicates the number of events in each matrix element. (B) Decoded information as a function of spatial granularity on real (white) and chance (grey) data for forward trials. (C) Decoding accuracy as a function of spatial granularity. (D) Decoding error as a function of the error position within the confusion matrix for forward trials. The color code indicates spatial granularity. In panels (A-D), data from 15 imaging sessions in 4 animals. (E-H) Same as in (A-D) for trials in the backward direction. Data from 17 imaging sessions in 4 animals. In (B, C, F, G) data are presented as mean ± s.e.m. See also Extended Data Table 5.

**Extended data Figure 8.**
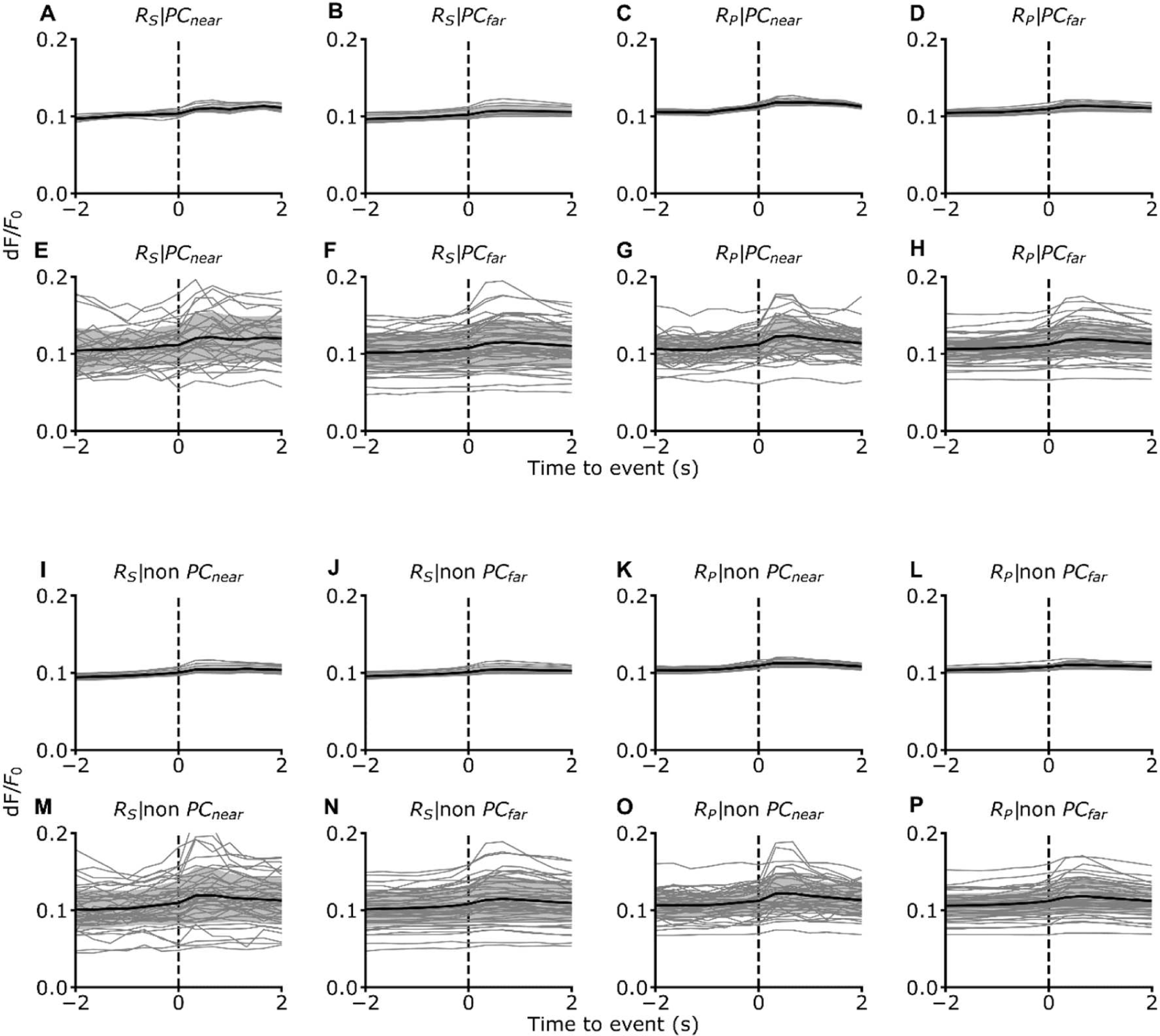
Temporal relationships between astrocytic and neuronal signals. (A-D) Event triggered average of astrocytic calcium responses. Calcium responses of putative receiver (R) ROIs are aligned to calcium events of neuronal place cells (PC). Astrocytic receiver ROIs could be in the soma (s) or processes (p). Neuronal cells were classified as being close (≤ 15µm) or far (> 15µm) from astrocytic receiver ROIs. Data from 11 imaging sessions in 7 animals. The black line indicates the mean, the shaded area the standard deviation. (E-F) Same as in (A-D) but for receiver ROIs belonging to the same astrocyte (N = 23 cells from 11 imaging sessions in 7 animals). (I-L) Same as in (A-D) but calcium responses of putative receiver (R) ROIs are aligned to calcium events of non-spatially informative cells (non PC). Data from 11 imaging sessions in 7 animals. (M-P) Same as in (I-L) but for receiver ROIs belonging to the same astrocyte (N = 48 astrocytes from 11 imaging sessions in 7 animals).

**Extended data Figure 9.**
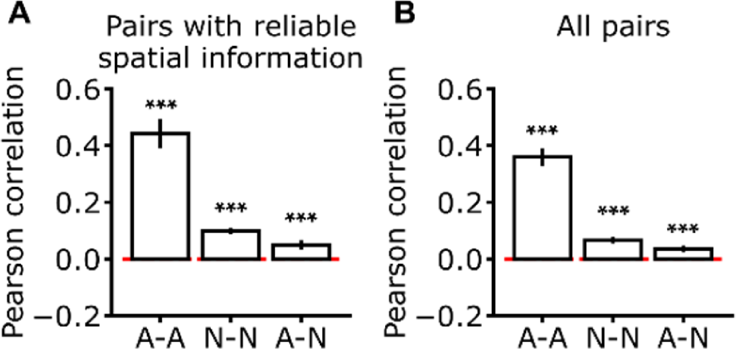
Pairwise correlations of calcium signals during virtual navigation. (A-B) Pearson correlation for different pairs of ROIs. Pairs were composed either of two astrocytic ROIs (A-A), two neuronal ROIs (N-N), or one astrocytic and one neuronal ROI (A-N). Red line indicates the zero correlation level. In (A), only results for ROI pairs with reliable spatial information are reported. In (B), results for all possible pairs are displayed. In (A), pairwise correlations of pairs with reliable spatial information are greater than 0, p = 2E-4, p = 7E-5, p = 3E-3, for A-A, N-N, and A-N pairs, respectively, Wilcoxon Rank sums test. In (B), pairwise correlations of all pairs are greater than 0, p = 7E-5, p = 7E-5, p = 1E-3, for A-A, N-N, and A-N pairs, respectively, Wilcoxon Rank sums test. Data are presented as mean ± s.e.m from 11 imaging sessions on 7 animals.

**Extended data Figure 10.**
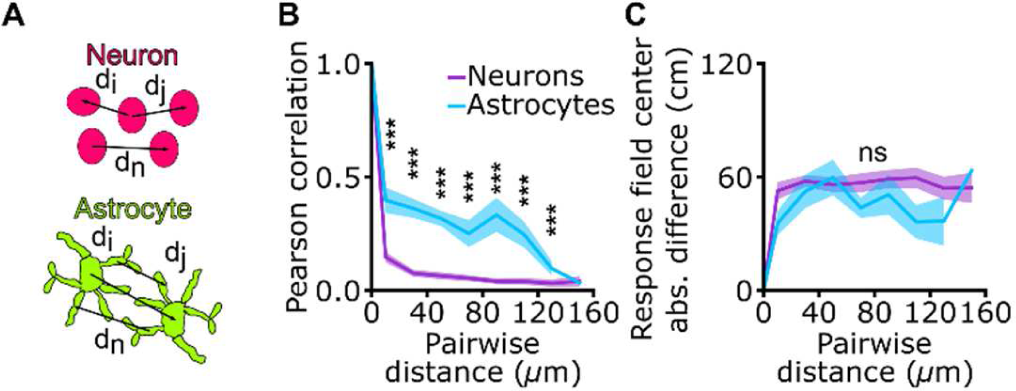
Pairwise correlation of calcium signals and difference in field position as a function of pairwise distance. (A) The distance (d) between the centers two ROIs comprising a pair is computed for all astrocytic (top) and neuronal (bottom) ROIs. (B-C) Pearson correlation (B) and difference between response field position (C) as a function of pairwise distance for pairs of astrocytic ROIs with reliable spatial information (cyan) and pairs of neuronal ROIs with reliable spatial information (purple). Data are expressed as mean ± s.e.m. from 11 imaging sessions on 7 animals. (A) p = 8E-4, p = 8E-4, p = 1E-4, p = 1E-3, p = 1E-3, p = 1E-3, p = 8E-4, and p = 2E-1 for 10, 30, 70, 90, 110, 130, and 150 µ m pairwise distances, respectively. Two-sample Kolmogorov-Smirnov test with Bonferroni post-hoc correction. (B) p = 1, p = 1, p = 0.7, p = 1, p = 1, p = 1, p = 0.2, and p = 0.2 for 10, 30, 70, 90, 110, 130, and 150 µ m pairwise distances, respectively. Two-sample Kolmogorov-Smirnov test with Bonferroni post-hoc correction.

**Extended data Figure 11.**
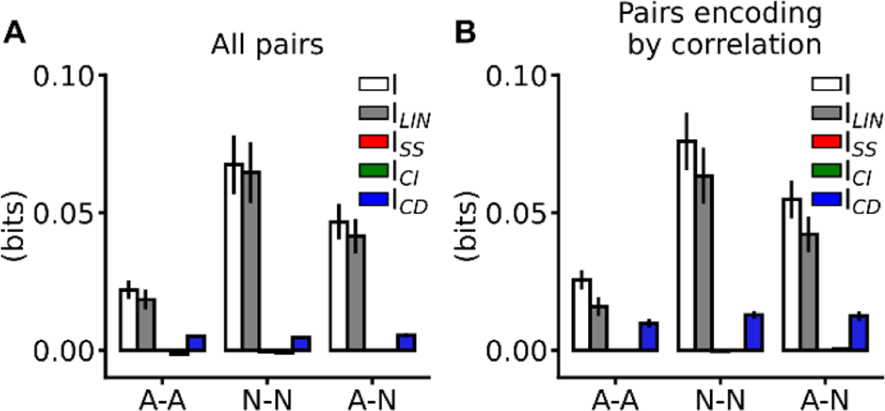
Position-dependent correlations contribute to synergistic information encoding. (A, B) Information breakdown for the different types of ROI pairs: two astrocytic ROIs (A-A), two neuronal ROIs (N-N), or one astrocytic and one neuronal ROI (A-N). Pairs were classified as synergistic (B) based on the value of ΔI (see Methods). I (white) is the mutual information about position encoded by the pair. I_LIN_ (grey) is the sum of the mutual information about position independently encoded in the response of each member of the pair. I_SS_ (red) is the redundant information component quantifying similarity in the responses of the members of the pair. I_CI_ (green) and I_CD_ (blue) quantify the information contribution of correlation independent or dependent on position, respectively. Data are represented as mean ± s.e.m and were collected in 11 imaging sessions on 7 animals.

**Extended data Figure 12.**
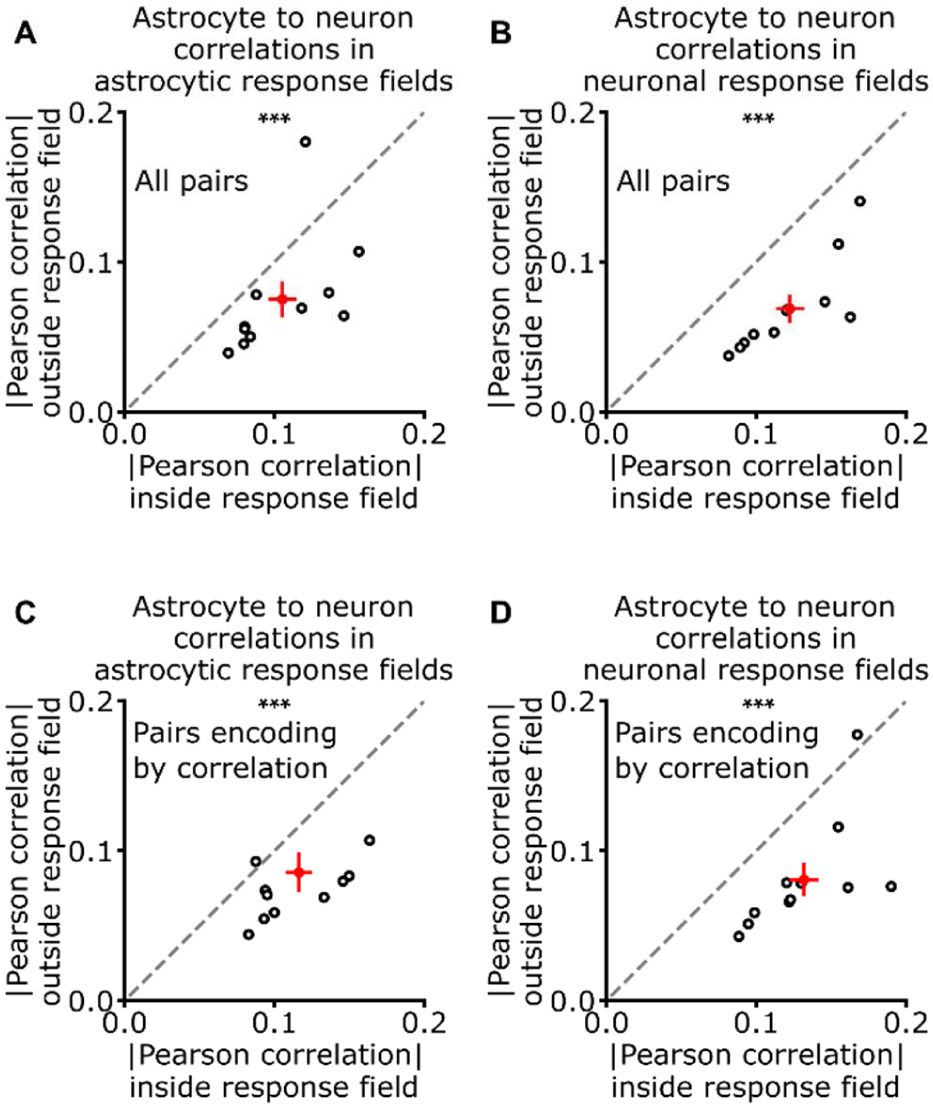
Correlation between astrocytes and neurons is animal’s position-dependent. (A-D) Scatterplot of the absolute value of Pearson correlation outside the response field against the absolute value of Pearson correlation inside the response field for pairs comprising one astrocytic and one neuronal ROI. Black open dots show averages of each imaging session, the red cross shows the mean ± s.e.m. (A, B) Correlations were measured for all possible pairs. In (A), correlations are computed with respect to astrocytic response field (mean correlation inside the response field 0.11 ± 0.01; mean correlation outside the response field 0.07 ± 0.01, p = 6.4E-3 Wilcoxon Rank sums test). In (B), correlations are computed with respect to neuronal response field (mean correlation inside the response field 0.12 ± 0.01; mean correlation outside the response field 0.07 ± 0.01, p = 1.1E-3 Wilcoxon Rank sums test). (C, D) Same as (A, B) but correlations were computed only on synergistic pairs based on the value of ΔI (see Methods, Fig. 5, and Extended Data Fig. 11). In (C), correlations are computed with respect to astrocytic response field (mean correlation inside the response field 0.12 ± 0.01; mean correlation outside the response field 0.09 ± 0.01, p = 7.8E-3 Wilcoxon Rank sums test). In (D), correlations are computed with respect to neuronal response field (mean correlation inside the response field 0.13 ± 0.01; mean correlation outside the response field 0.08 ± 0.01, p = 1.8E-3 Wilcoxon Rank sums test). For each pair of ROIs, correlations were computed averaging 100 resampling to compensate unbalanced observations inside and outside the response field. Data from 11 imaging sessions on 7 animals.

**Extended data Fig 13.**
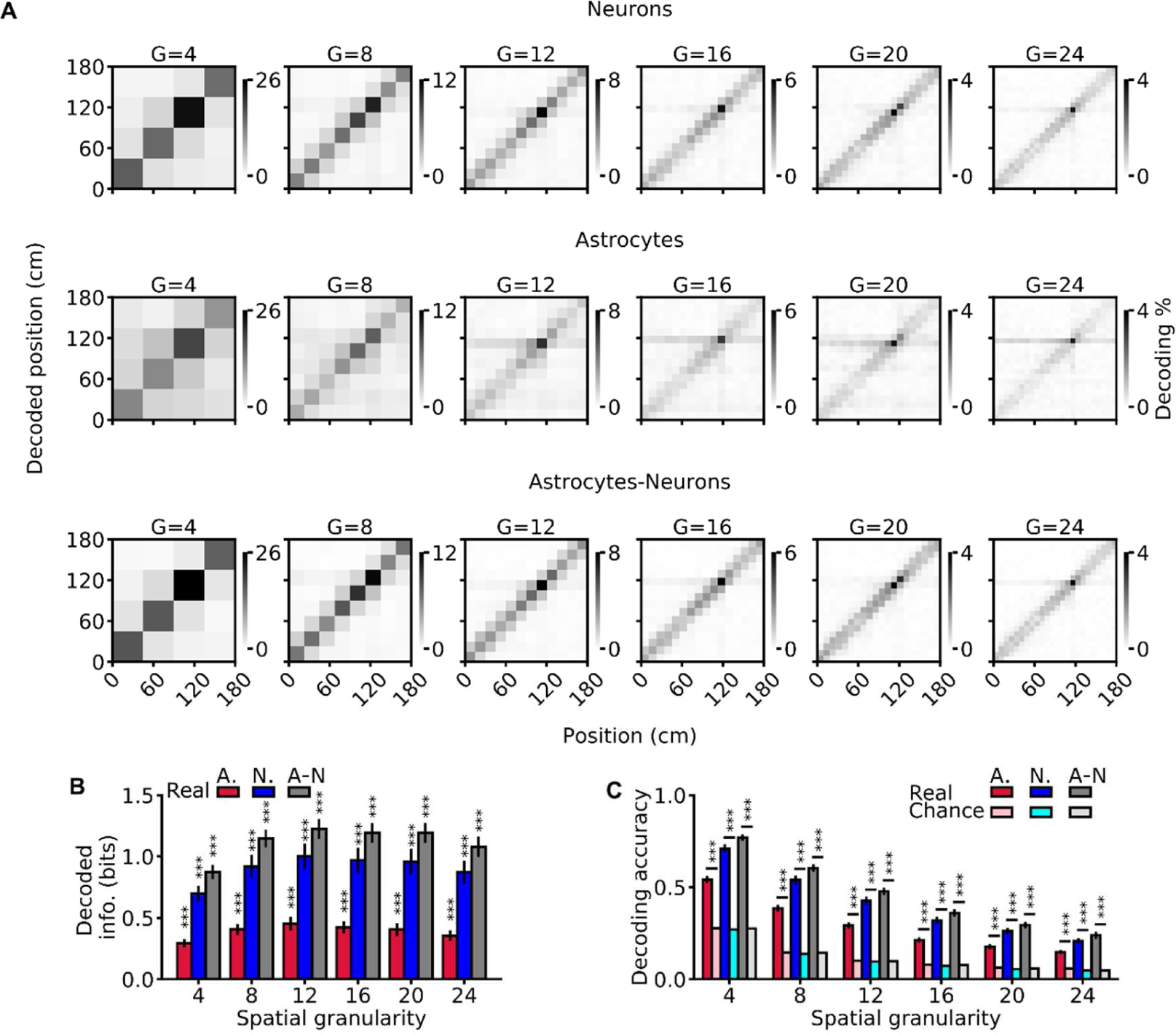
Decoding the animal’s position from neuronal and astrocytic population vectors. (A) Confusion matrices of a SVM classifier decoding the mouse’s position using population vectors comprising neuronal (top), astrocytic (middle), and neuronal + astrocytic ROIs (bottom) for various spatial granularities (G = 4, 8, 12, 16, 20, 24). The true position of the animal is shown on the x-axis and the decoded position on the y-axis. Grey scale indicates the percentage of occurrence of each matrix element. (B) Decoded mutual information between predicted and real position in the linear track and (C) decoding accuracy for the different population vectors as a function of spatial granularity. In B and C, asterisks indicate significance against chance level (Extended Data Table 4 and Table 6). Data are displayed as mean ± s.e.m and were collected in 11 imaging sessions from 7 animals.

**Extended data Table 1.**
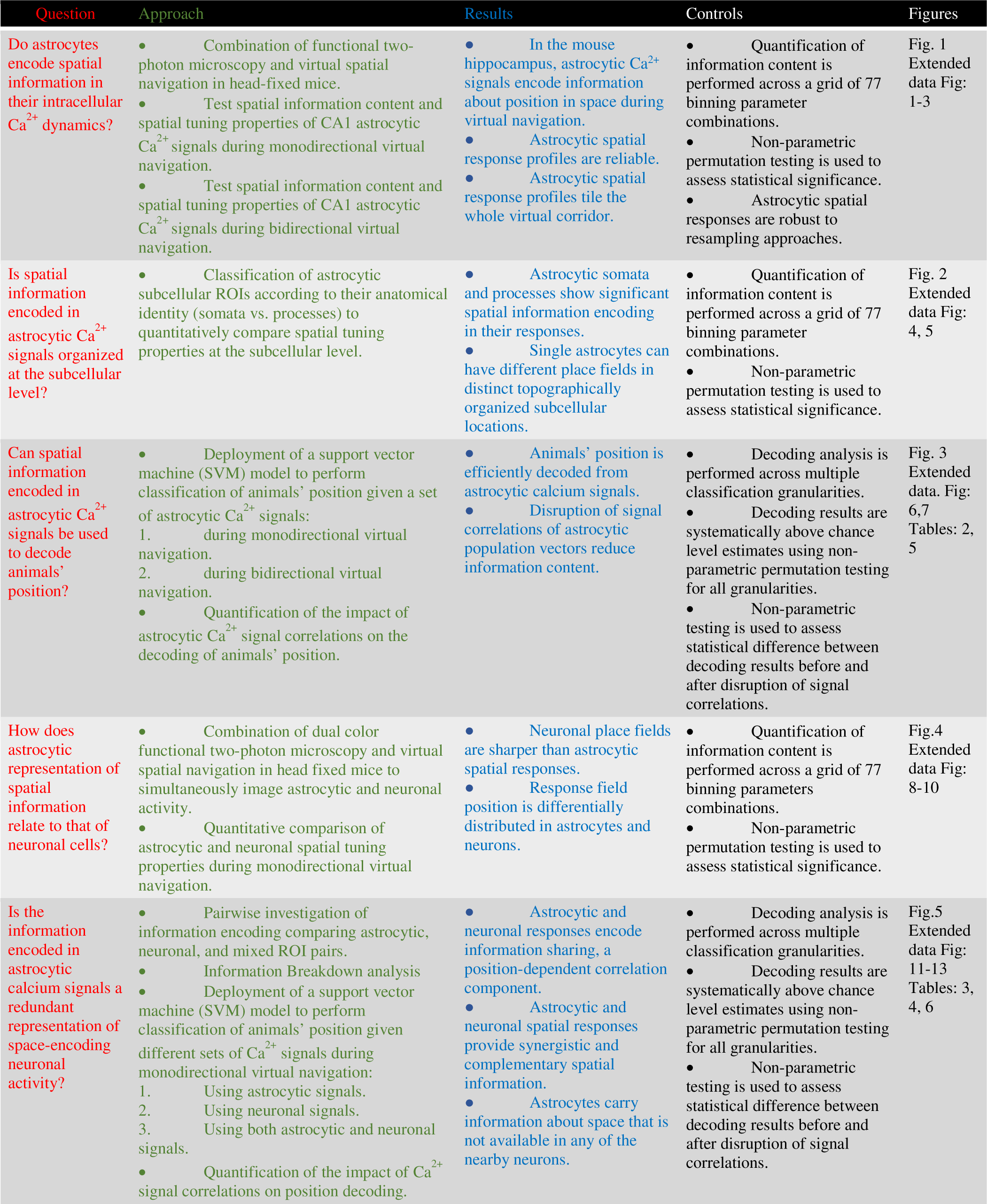
Outline and summary of experiments.

**Extended data Table 2.**
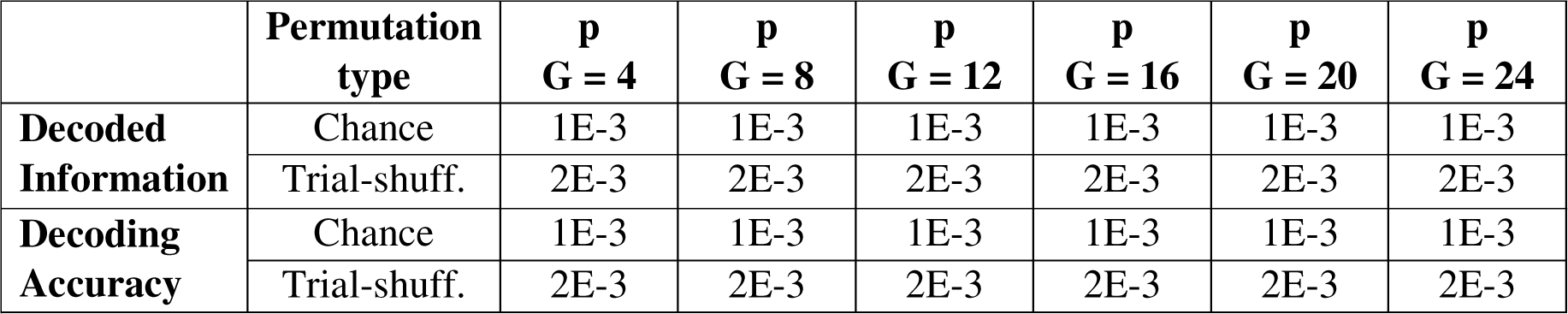
Hypothesis testing: decoding performance about animal’s spatial location from astrocytic calcium signals during monodirectional virtual navigation. p-values for one-tailed non-parametric permutation tests as a function of decoding granularity for decoded information (see Fig. 3B) and decoding accuracy (Extended Data Fig. 6). For each imaging session and each granularity, null distributions were obtained with 1000 and 500 iterations to estimate chance level and trial-shuffling, respectively (see Methods). Data from 7 imaging sessions from 3 animals.

**Extended data Table 3.**
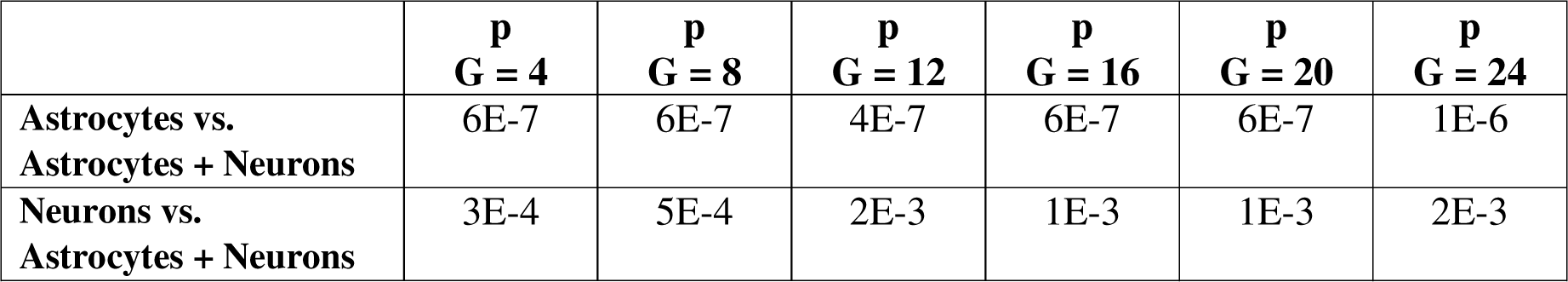
Comparison of decoding information about animal’s spatial location from neuronal and astrocytic population vectors. p-values for two-tailed paired t-tests with Bonferroni-correction for decoded information of animal’s spatial location from population vectors comprising all astrocytic ROIs *vs* all ROIs of both types (top row) and all neuronal ROIs *vs* all ROIs of both types (bottom row) during monodirectional virtual navigation shown in Fig. 5. Data from 11 imaging sessions from 7 animals.

**Extended data Table 4.**
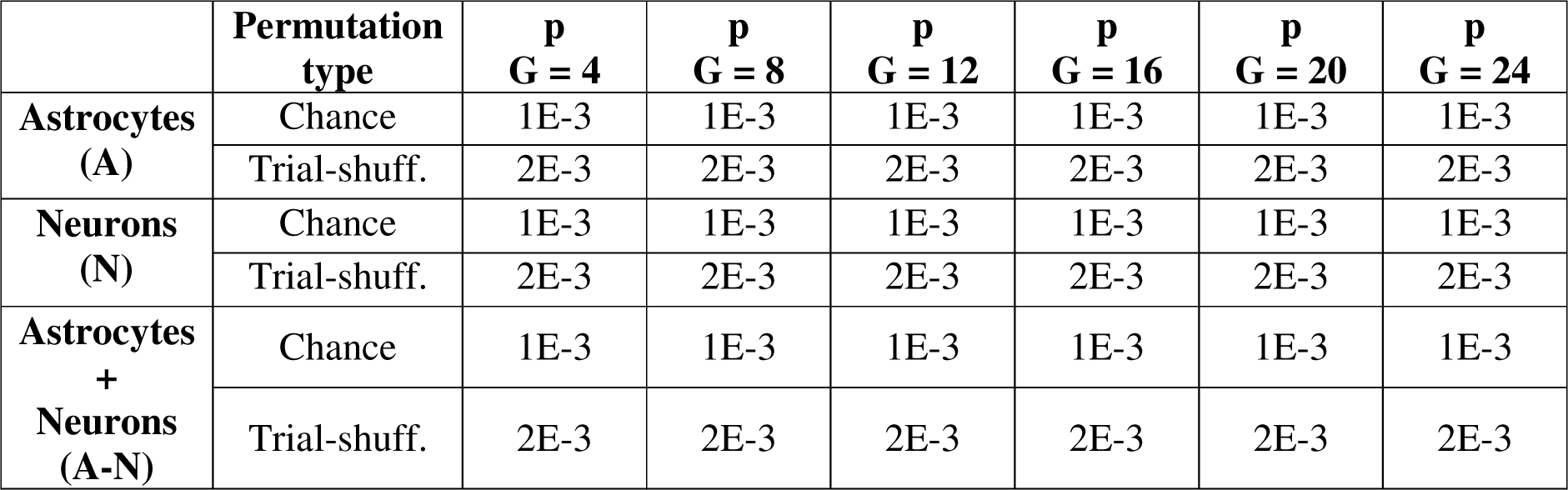
Hypothesis testing: decoding information about animal’s spatial location from neuronal and astrocytic population vectors. p-values for one-tailed non-parametric permutation tests for decoding information from population vectors comprising either all astrocytic (top row), all neuronal (middle row), or ROIs of both types (bottom row) during monodirectional virtual navigation (see Fig. 5 and Extended Data Fig. 13). Significance levels are reported as a function of decoding granularity. For each imaging session and each granularity, null distributions were obtained with 1000 and 500 iterations to estimate chance level and trial shuffling, respectively (Methods). Data from 11 imaging sessions from 7 animals.

**Extended data Table 5.**
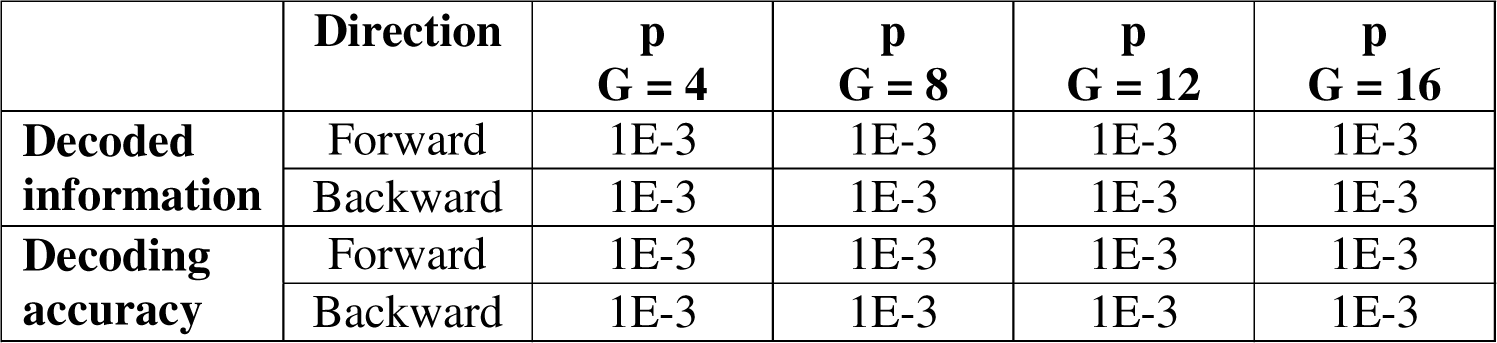
Hypothesis testing: decoding performances about animal’s spatial location from astrocytic calcium signals during bidirectional virtual navigation. p-values for one-tailed non-parametric permutation tests as a function of decoding granularity for decoded information (see Extended Data Fig. 7B, F) and decoding accuracy (see Extended Data Fig. 7C, G). Decoding performance is reported for forward- and backward-running directions (see Extended Data Fig. 7). For each imaging session and each granularity, null distributions were obtained with 1000 iterations to estimate chance level (Methods). Data from 15 imaging sessions in 4 animals for forward-running direction. Data from 17 imaging sessions in 4 animals for backward-running direction.

**Extended data Table 6.**
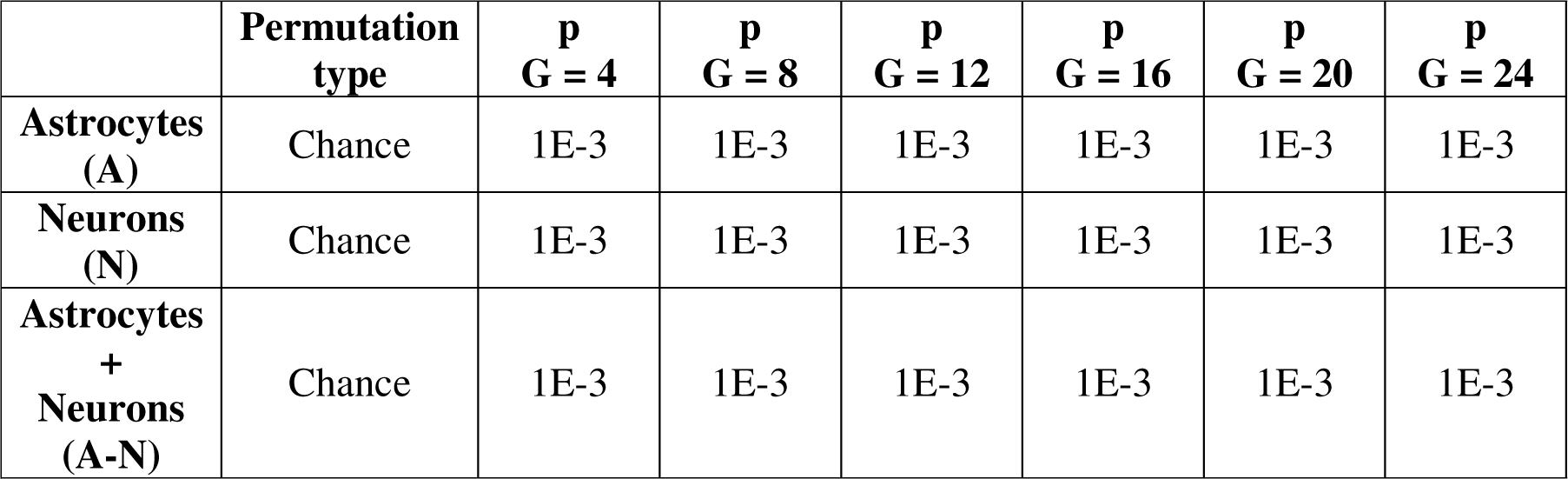
Hypothesis testing: decoding accuracy about animal’s spatial location from neuronal and astrocytic population vectors. p-values for one-tailed non-parametric permutation tests for decoding accuracy from population vectors comprising either all astrocytic (top row), all neuronal (middle row), or all ROIs of both types (bottom row) during monodirectional virtual navigation (see Extended Data Fig. 13). Significance levels are reported as a function of decoding granularity. For each imaging session and each granularity, null distributions were obtained with 1000 iterations to estimate chance level (Methods). Data from 11 imaging sessions from 7 animals.

